# From Recovery to Risk: Intensifying ENSO Events Drive Extirpation Risk in a Marine Predator

**DOI:** 10.64898/2026.10.01.754291

**Authors:** Haze W. Y. Chung, Susana Cárdenas-Alayza, Joanna Alfaro-Shigueto, Kevin Farfán-Hernandez, María José Ganoza-Gallardo, Jeffrey C. Mangel, Katrina J. Davis, Michael B. Bonsall

## Abstract

Climate change poses a major threat to marine mammal populations, many of which remain vulnerable after years of overexploitation during the 17th to 20th centuries. Extreme climate events have grown in both frequency and severity in recent decades, and this trend is projected to accelerate under continued greenhouse gas emissions. Here, we assess how this intensification affects a top marine predator at the epicentre of extreme El Niño events – the Peruvian population of *Otaria byronia* (South American sea lion). We first apply a Bayesian population model that best predicts observed population dynamics to reconstruct female *O. byronia* population trends, estimate vital rates, and evaluate the effects of environmental drivers on survival and reproduction. We then use the same model to project extirpation risk under multiple future climate scenarios reflecting anticipated increases in both the intensity and frequency of extreme El Niño events. Our reconstruction reveals that the Peruvian *O. byronia* population has shown considerable resilience to episodic El Niño events over the past quarter-century, likely sustained by high female survival during normal and favourable years. However, under a scenario of moderate increases in El Niño frequency and intensity, the probability of extirpation over the next 50 years is 83.4% higher than under current conditions. As disturbance regimes intensify globally, we suggest that resilience-oriented management frameworks, grounded in species-specific demographic data and environmental context, as demonstrated here, will be essential for safeguarding marine mammal populations against an increasingly volatile climate.

## Introduction

Climate change poses a major risk to marine mammal populations, alongside operational interactions with fisheries (Avila et al., 2018; Kovacs et al., 2012; Simmonds & Issac, 2007). Key forms of climate-related threats include the loss of sea ice for ice-associated species (Laidre et al., 2015; Ouled-Cheikh et al., 2024; Stenson & Hammill, 2014), shifts in prey availability and nutritional quality (Bas et al., 2025; Germishuizen et al, 2024; Harvey & Moore, 2017; Hunt Jr et al., 2002; Kortsch et al., 2015), and extreme climate events that trigger mass mortality (de Oliveira, 2012; Laake et al., 2018). However, before global surface temperatures began to accelerate in the 1970s (Hansen et al., 2010), marine mammal abundances were already severely reduced by human exploitation between the 17th and 20th centuries. During this period, whale populations are estimated to have reduced by ∼82% relative to historical baselines, and other marine mammal species by 96%, with the lowest levels observed in the 1900s (Lotze & Worm, 2009). Since then, fewer than half of marine mammal populations have shown significant increases in abundance, while many others exhibit declining or uncertain population trends (Magera et al., 2013). Monitoring marine mammal populations is the first step in assessing climate-related risks and developing strategies to mitigate further declines.

Being situated in the tropical Eastern Pacific, Peru is one of the countries most affected by extreme climate events associated with the El Niño–Southern Oscillation (ENSO). While ENSO events occur every 2–7 years, extreme episodes are less frequent and tend to occur on a decadal scale (Racault et al., 2017). El Niño events are the warm phase of ENSO and occur when trade wind patterns shift, pushing warm, nutrient-poor waters across the Pacific. This reduces nutrient availability and leads to elevated sea surface temperatures (SST) and lower primary productivity (Racault et al., 2017). Nonetheless, the tropical Eastern Pacific is one of the world’s most productive marine ecosystems, sustained by intense upwelling associated with the northern Humboldt Current System (Bakun & Weeks, 2008; Ramos et al., 2022). Although several Eastern Boundary Upwelling Systems share comparable physical characteristics, including the California and Benguela systems, fishery yields in the Humboldt region are remarkably higher (Chavez & Messié, 2009). The region supports the world’s largest single-species fishery, which at its peak produced 20% of the global fish catch from an area covering less than 0.01% of the global ocean (Chavez & Messié, 2009). Benefiting from abundant prey sources, at least 29 marine mammal species have been recorded in the region, including fur seals, sea lions, whales, dolphins, and the marine otter (Majluf & Reyes, 1989). Among these species, the most conspicuous are the two pinnipeds, South American sea lions (*Otaria byronia*) and South American fur seals (*Arctocephalus australis*), with colonies distributed along the entire Peruvian coastline. *O. byronia* is the larger and more numerous of the two, and today represents the dominant pinniped species in Peru (Cárdenas-Alayza et al., 2021).

*O. byronia* is currently listed as Least Concern on the IUCN Red List (Cárdenas-Alayza et al., 2016), though the Peruvian population is particularly vulnerable to the significant climate fluctuations caused by ENSO (Cárdenas-Alayza et al., 2021; de Oliveira et al., 2012). For instance, the 1997–1998 El Niño event resulted in an 81% decline in observed *O. byronia* abundance (Arias-Schreiber & Rivas, 1998). The onset of El Niño creates complex ecosystem changes, but generally leads to a reduction of coastal productivity that cascades through the bottom-up Peruvian ecosystem (Lehodey et al., 2020; Taylor et al., 2008). Previous studies have documented the adverse effects of reduced anchovy biomass, a key prey species of *O. byronia*, especially in El Niño years (Ramos et al., 2022; Soto et al., 2006). Prey shortages can increase foraging effort in pinnipeds, reducing maternal attendance and elevating pup mortality (McMahon & Burton, 2005; Soto et al., 2006; Speakman et al., 2020). Extreme climate events affecting the *O. byronia* population have intensified in both frequency and severity in recent decades (Lee & McPhaden, 2010; Nicholls, 2008). Recent analyses indicate a ∼25% increase in ENSO variability since the pre-industrial era, suggesting a link to anthropogenic greenhouse forcing (Cai et al., 2021). This trend is projected to continue, with the frequency of extreme El Niño events expected to rise by 59%, from 5.57□±□0.81 during 1900–1999 to 8.81□±□1.03 over 2000–2099 (Cai et al., 2021). Given these escalating pressures, conserving *O. byronia* into the future requires assessing the extirpation risk under future climate scenarios.

A persistent challenge in predicting pinniped population responses to future climate change is accurately estimating past and present population dynamics. This difficulty arises because only a proportion of individuals are present on land during censuses. Inferring the total population size from this “haul-out” proportion requires additional tagging or telemetry data (Grandi et al., 2021; Hammond et al., 2021; Sharples et al., 2009). Some studies also use pup counts with a pup correction factor (also called a pup multiplier) to estimate total abundance (Berkson & DeMaster, 1985; Robertson & Chilvers, 2011; Wood et al., 2022). This approach assumes that pups, which remain on land for extended periods during the breeding season, can be reliably counted (Trites & Larkin, 1996; Hammond et al., 2021). However, several studies have noted potential bias introduced by pup correction factors, for example, when the population is not at a stable age distribution (Berkson & DeMaster, 1985; Karamanlidis, 2024; Laake et al., 2018). Other common challenges in ecological censuses, including yearly environmental stochasticity, survey errors, and incomplete counts, further complicate analyses of population dynamics by introducing uncertainty and potential bias into estimates of abundance (Conn et al., 2015; Schaub & Kéry, 2021).

A promising way to overcome these limitations is a Bayesian state-space model, which explicitly separates observation error from underlying population processes, thereby enabling the estimation of haul-out proportions. Variants of this modelling approach have been applied across a wide range of taxa, including sea otters, lions, and birds (Bauer et al., 2015; Schaub & Kéry, 2021; Tinker et al., 2021). The Bayesian framework allows the integration of prior knowledge with long-term count data, facilitating the imputation of missing observations and the estimation of key demographic parameters. Its flexibility also enables the incorporation of different drivers of population dynamics, as demonstrated in harbour seal studies, such as the negative effects of predator abundance on pup survival (Caillat et al., 2019) and impacts of extreme SST anomalies on Guadalupe fur seals (Juárez-Ruiz et al., 2022). This framework is therefore particularly appropriate for assessments of the population dynamics of *O. byronia* in Peru, given the need to assess the effects of extreme El Niño events and the scarcity of vital rate estimates and haul-out probabilities for this species.

This study assesses the extirpation risk (a ≥80% decline over three generations) of Peruvian *O. byronia* under future climate change by quantifying the impact of environmental variability on population dynamics. We utilized stage-based count data in Peru (1997-2022) to parameterize a deterministic, female-only population model, estimating vital rates (survival and birth rates) and total female abundance. We then embed the population model within a state-space framework to account for unknown haul-out probability, environmental error, observation error and missing data (Aarts et al., 2019). We included three covariates influenced by ENSO-driven variability, which are SST, zooplankton concentrations, and anchovy biomass, to represent successive trophic levels, from physical to biological drivers. To identify the covariate that best explains population dynamics, we compared a base population model and three covariate-driven models. Focusing on the Humboldt Current System along the Peruvian coast, we apply the model that best predicts observed dynamics to reconstruct *O. byronia* population trends, estimate species-specific vital rates, and evaluate the effects of each covariate on survival and reproduction. Finally, we use the same model to project extirpation risk under multiple future climate scenarios that reflect anticipated increases in both the intensity and frequency of extreme El Niño events. Together, this framework identifies the mechanisms linking environmental variability to population viability and provides a scientific basis for conservation strategies that account for both ecological complexity and climate uncertainty.

## Methods

Below, we outline the procedure used to assess how environmental variability influences the Peruvian *O. byronia* population dynamics and to predict extirpation risk under future climate change. We first provide an overview of the data, modelling framework, covariates, and Bayesian implementation and evaluation. Finally, we present the sensitivity analyses and extirpation risk projections.

### 1. Study area and data sources

*O. byronia* inhabits a wide coastal range extending along the shores of South America, from Peru in the west to Brazil in the east. This species exhibits pronounced sexual dimorphism, with males reaching up to 3 meters and weighing between 300–350 kg, while females are smaller, averaging 2 meters and weighing approximately 150 kg (Cappozzo & Perrin, 2009). Females typically produce a single pup annually and reach sexual maturity at around 4.8 years (±0.5), when half of the population is reproductively active (Grandi et al., 2010). Pups are born onshore and remain near their mothers after birth.

In Peru, seventy of the 86 registered colonies are managed by the National Service of Natural Protected Areas by the State (SERNANP), while the remaining 16 unprotected sites host only 4% of the total population (IMARPE, 2020, Nr 149 and 070-2020-IMARPE/OGA). Most protected colonies are located within the Guano Islands, Islets, and Capes National Reserve System (RNIPG) and the Paracas National Reserve (RNP) (see Figure 1). Enforcement levels, ranger presence, and regulations on resource use vary considerably across these reserves; some sites are actively patrolled, whereas others are less supervised (Laínez del Pozo & Jones, 2021).

**Figure 1.**
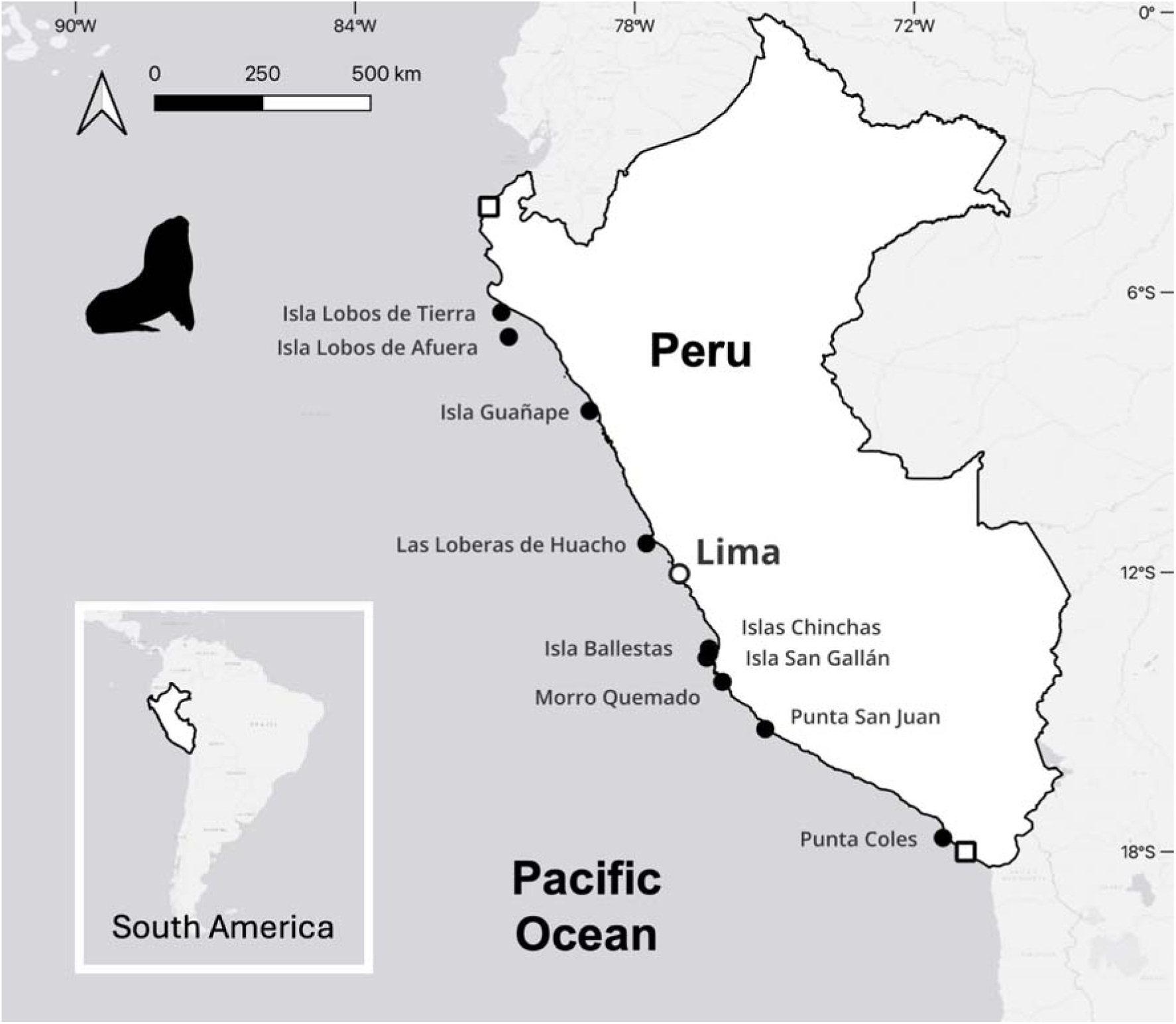
Coastline of Peru indicating the ten largest *O. byronia* colonies (black circles) reported in recent IMARPE reports (IMARPE, 2020–2022). White squares mark the start/end of the survey range along the coast. Figure created in QGIS 3.42.1 using the ESRI Gray basemap and shapefiles from the GADM database (www.gadm.org, version 2.5, July 2015).

We collated survey data collected by the Instituto del Mar del Perú (IMARPE) from 1997 to 2022. IMARPE has conducted annual surveys of *O. byronia* during the breeding season using binoculars from the shore or boat since 1997. The surveys cover all major colonies, from Los Órganos (4°10′ S) to Morro Sama (18°00′ S) (Figure 1); however, the number of recorded colonies varies across years due to changes in distribution. Surveys were conducted in all years except 2001 and typically lasted 2–3 weeks (IMARPE, 1997–2022). We defined the usual survey months, February and March, as the “survey period.” Our data came from two main sources: counts obtained through a request for information through the transparency platform (https://www.gob.pe/20399-solicitar-acceso-a-la-informacion-publica?child=20744) and population counts published in annual reports (IMARPE, 1997–2022). We excluded certain data points due to substantial survey delays, incomplete spatial coverage, or discrepancies between data sources; detailed retention criteria are provided in Supplementary Data SD1.

To account for *O. byronia* at sea, we estimated pup and adult haul-out probabilities using two additional datasets. We defined the pup haul-out ratio as the proportion of pups present on a randomly selected sampling date relative to the maximum number observed on any day during the survey period. The adult haul-out ratio was defined as the average proportion of adults present on a randomly selected sampling date (see SM3 for the rationale behind these definitions). The datasets were collected by the Punta San Juan Program, a long-term scientific field project that conducts weekly counts of pinnipeds thanks to agreements with government officials and SERNANP authorizations (https://www.puntasanjuan.org/english.php?lang=us). For adult females, we used telemetry data from four tagged individuals to estimate the proportion of time present in the rookery (Data collected under: RJ N° 019-2016-SERNANP-RNSIIPG). The data were collected in 2017, a neutral ENSO year, to avoid the influence of extreme conditions on haul-out behavior. We included only daytime presence-absence data (6□am to 6□pm) at the Punta San Juan rookery. The period used was February 26 to March 31, 2017, which overlaps with the IMARPE survey period. For pups, we used a long-term stage count time series from 2012 to 2024 (Data collected under specific interinstitutional collaboration agreements between Universidad Peruana Cayetano Heredia and SERNANP 2011-2016, 2017-2022 and RJ No. 013-2022-SERNANP-RNSIIPG). Observers conducted weekly land-based counts using 10×50 binoculars between 6 am and 9□am at accessible sites. These local, high-resolution data provided valuable information to improve estimates of stage-specific haul-out behavior in our population model.

### 2. State-space model structure

To reconstruct the pronounced interannual fluctuations in *O. byronia* populations, we developed a state-space model consisting of a process and an observation component. We implemented the model using a Bayesian framework in Stan (Stan Development Team, 2023), which employs Hamiltonian Monte Carlo (HMC) sampling to enable efficient posterior inference. The following sections describe each component in detail; prior specifications are provided in Supplementary Methods SM3.1.

#### 2.1. The process model

The process model is a female-only stage-based population model spanning 1997 to 2022 that consists of five stages: newborn pups, juveniles in their first through third years, and reproductive adults (Figure 2). This discretisation reflects biological transitions, as only adults (age 4 and older) are capable of reproduction, while younger age groups have lower survival (Grandi et al., 2016). Juveniles were separated into three substages (e.g., age 1, age 2, and age 3) to match the model’s annual time step. This structure is consistent with stage-based approaches in pinniped demography (Caillat et al., 2019; Davis, 2022). We opted for a simplified female-only population model and treated the Peruvian population as demographically closed (see Supplementary Methods SM1). The largest population size of *O. byronia* was observed in the 1997 survey, we therefore assumed the initial 1997 population structure followed the stable-stage distribution implied by the priors of the vital rates.

**Figure 2.**
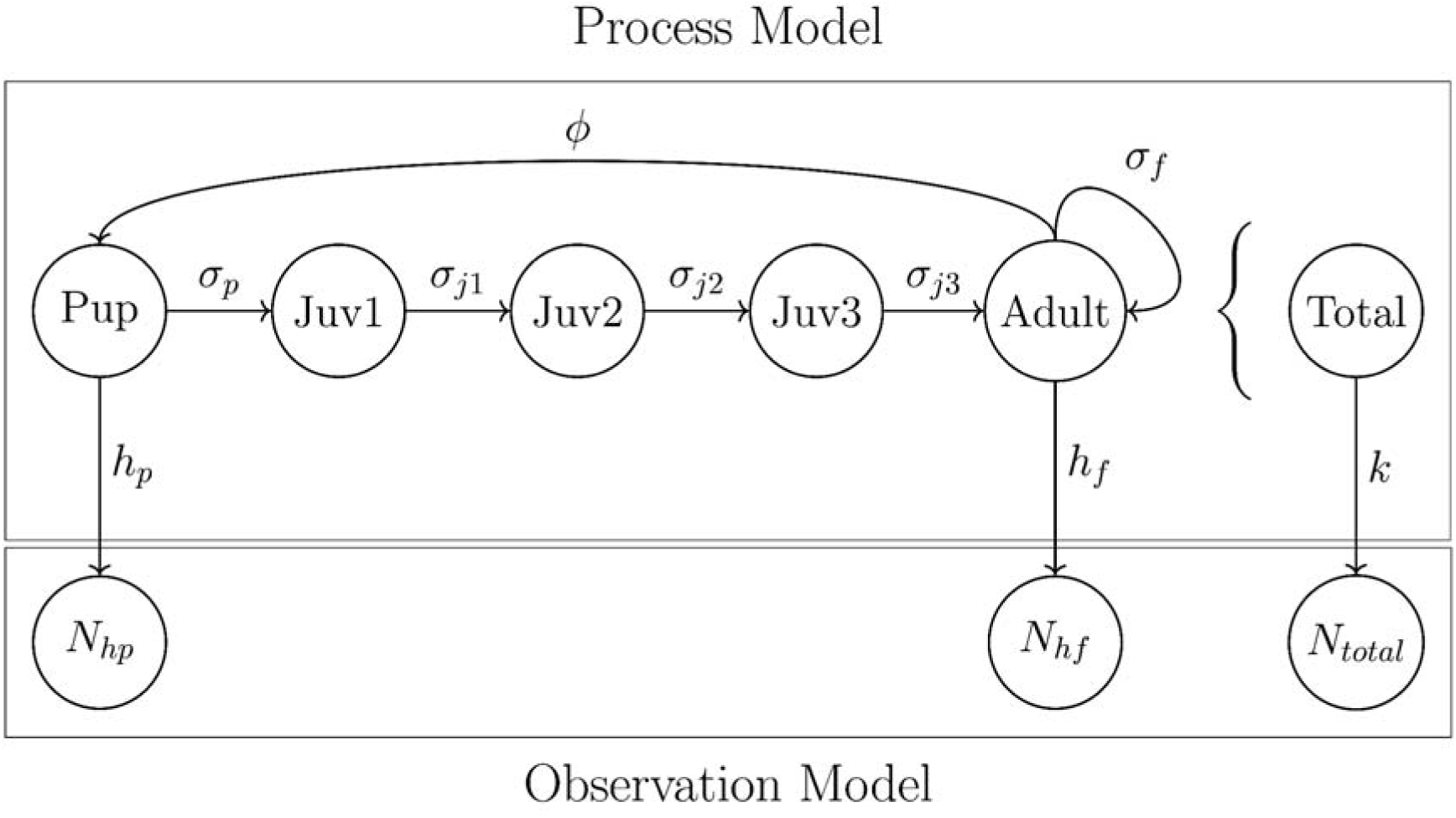
Schematic representation of the state-space model used to assess the effect of extreme El Niño events on the Peruvian population of *O. byronia*, which comprises a process model and an observation model. The process model outlines the life cycle of the female *O. byronia* population across five stages: pup, juvenile (ages 1-3) and adult. The arrows in the process model indicate the progression through these stages. We denote survival by and birth rate by, where, (for ages 1–3), and represent pup, juvenile, and adult survival, respectively. The observation model links latent state variables to three data streams: pup haul-out counts ( ) and female adult haul-out counts ( ), scaled by fixed haul-out proportions and respectively, and total counts (both sexes,) approximated from the latent female population via a time-varying scaling factor (defined in Methods section 2.2). Each data stream enters a negative binomial likelihood with a separate overdispersion parameter.

The key ecological parameters that we estimated in the process model were birth rate (□), pup survival (σ_p_), three juvenile survival rates (σ_ji_) and adult survival (σ_f_). Each time step in the model corresponds to the period (i.e., year) between consecutive breeding seasons (February to March). Birth rate (□) refers to the reproduction probability of female adults, assuming each adult female produces one pup per reproductive event and that offspring are evenly split between sexes (Campagna & Boeuf, 1988; Soto et al., 2004). To model the effect of covariates on the survival and birth probabilities, we used a hazard-based discrete-time formulation with a complementary log–log (cloglog) transformation and quadratic terms to capture asymmetric, non-linear biological responses under extreme environmental conditions (Prentice & Gloeckler, 1978; Tinker et al., 2021; Alam & Linero, 2025). These vital rates were modeled as functions of the relevant covariate value at each time step, *Cov_t_*, which represents a summary of environmental conditions over a specified temporal window (Figure 3). To assess the appropriateness of our formulation, we conducted a set of sensitivity analyses comparing the best model’s performance to alternative specifications (see section 5 of Methods).

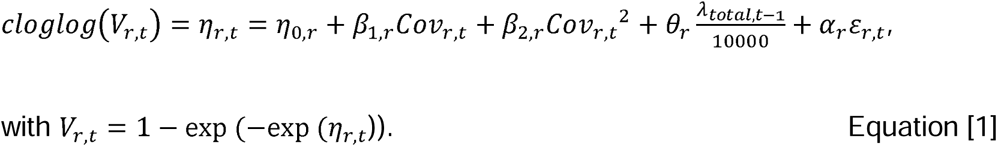

**Figure 3.**
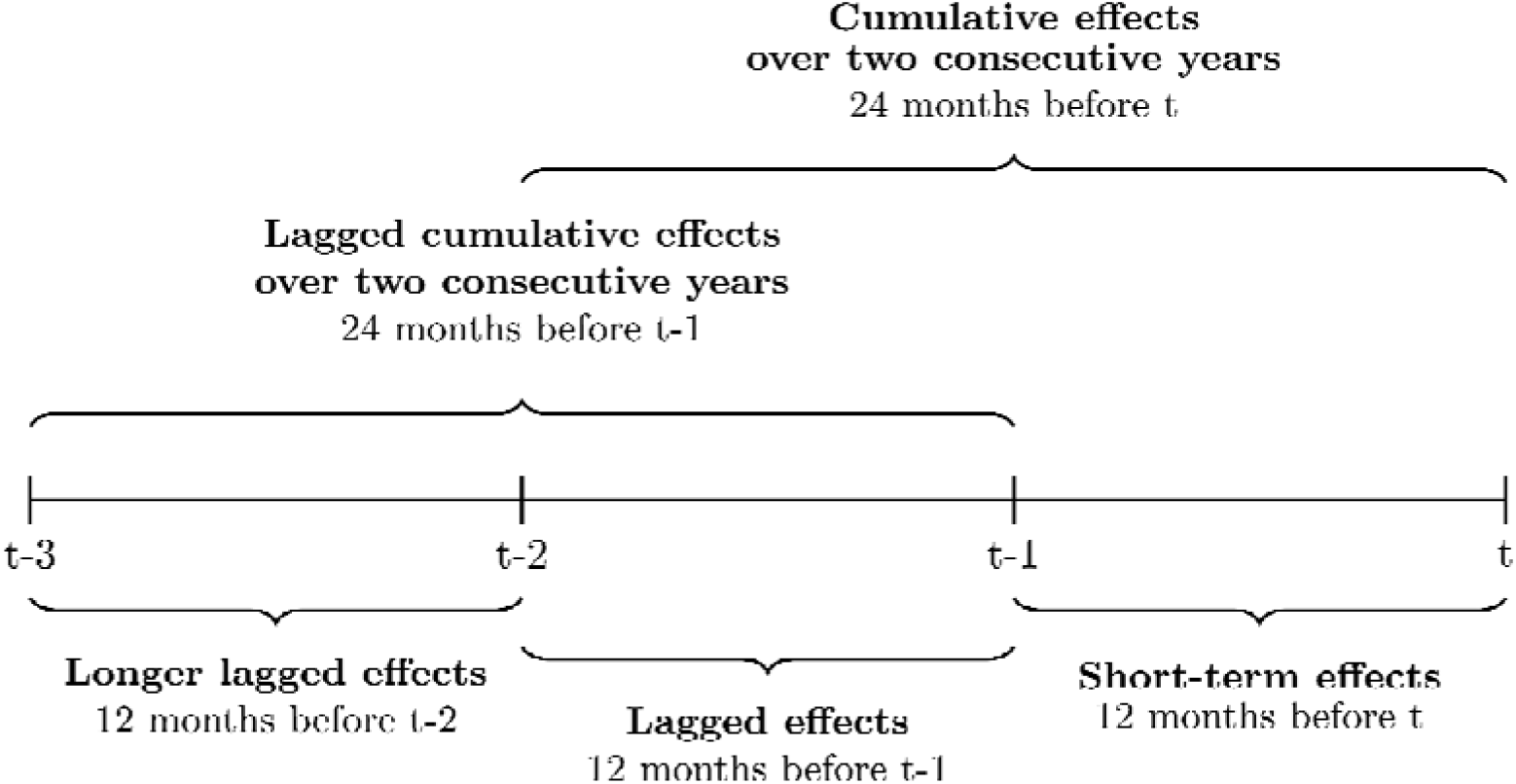
Temporal windows used to evaluate environmental effects, including sea surface temperature, zooplankton concentration, and anchovy biomass, on the population dynamics of *O. byronia* in Peru.

In equation [1], *v_r,t_* denotes the vital rate *r* at time *t*, including the stage-specific survival rate and birth rate. The linear predictor *η_r,t_* is defined on the cloglog scale and aggregates baseline demography, environmental covariate effects, density dependence, and stochastic variation. The baseline intercept *η_0,r_* is parameterised via a baseline probability *γ_r_* ∊ (0,1) as *η_0,r_*= log(– log(1 – *γ_r_*)), such that *γ_r_* represents the expected vital rate when covariate and stochastic effects are zero. The coefficients *β*_1,*r*_ and *β*_2,*r*_ describe linear and quadratic effects of the environmental covariate, respectively. The contribution of environmental stochasticity is scaled by *α_r_*, where *ε_r,t_*∼*Normal*(0,1) is a shared annual perturbation across all vital rates. Density dependence is incorporated through the total female population size in the previous year, *λ_total,t_*_—1_, scaled by 10,000 for numerical stability. The coefficient *θ_r_* follows standard formulations of lagged density feedback in discrete-time population models (Caswell, 2001). In the base model, covariate effects (*β*_1,*r*_ and *β*_2,*r*_) are omitted.

#### 2.2. The observation model

The observation model included three components, each contributing to the likelihood by comparing observed counts to estimated abundances. We used the negative binomial distribution to account for overdispersion in the count data due to variability in detection, animal behavior, and environmental conditions (Schaub & Kéry, 2021; Stoklosa et al., 2022). The first and second components are based on adult female counts (n = 13) and pup counts (n = 16). We focused on these two groups due to their higher use of breeding sites. The haul-out abundances are obtained by multiplying the estimated number of adult females and pups by their respective haul-out ratios, *h_f_* and *h_p_*. These specific haul-out ratios were estimated within the model and assumed to remain constant across all years.

The third component of the likelihood is calculated using total counts (including both sexes, n = 19). The strong 1997–1998 El Niño event is known to have greatly reduced birth rates (Soto et al., 2004). However, female adult and pup counts are not available before 2000. To incorporate earlier population trends, we included total counts in the observation model. Total counts were linked to the latent female population estimate, *λ_total_*, with the yearly expected total count of both sexes estimated by *λ_total,t_δ* exp(*α_u_ε_u,t_*). The parameter *δ* is a scaling factor and *ε_u,t_*∼*Normal*(0, 1) is an annual perturbation scaled by *α_u_*, together constituting the time-varying scaling factor *k* in Figure 2. We estimated separate dispersion parameters for each of the three data types in the negative binomial likelihood. We treated missing counts for specific years as latent variables and inferred them from the stochastic population trajectory.

Lastly, since pup multipliers are commonly used in pinniped monitoring (e.g., Wood et al., 2022), we evaluated whether total observed individuals or observed pups better serve as proxies for true abundance by comparing their consistency and predictive utility. After estimating the total females, we computed correction multipliers as the ratio of the estimated total females to observed total or pup counts of both sexes. We modeled them as smooth functions of estimated total female abundance using Generalized Additive Models (GAMs) with a low-complexity smoothing term (k = 3; see Chung et al., 2026).

### 3. Covariates

To evaluate the influence of environmental drivers on population dynamics, three covariates were included in the analysis: SST, zooplankton concentration, and anchovy biomass. Environmental conditions in one year can influence survival or reproduction in subsequent years due to delayed physiological responses or changes in individual condition (Harrison et al., 2011). We therefore tested all pairwise combinations of the five temporal averaging windows for birth rate and survival respectively, where data permitted (Figure 3). This approach resulted in a total of 59 models for the full time series spanning 1997–2022 (25 models each for SST and zooplankton concentrations, and 9 for anchovy biomass due to data constraints). The data source for each covariate and the adjustments made to extract their respective time series are listed in Supplementary Methods SM2.

### 4. Model convergence and assessment

We estimated model parameters for all analyses using Bayesian inference implemented in R version 4.4.1 (R Development Core Team, 2023), Stan 2.32.0 (Stan Development Team, 2023), and CmdStanR 2.36.0 (Gabry et al., 2025). We ran seven concurrent MCMC chains with at least 12,000 iterations each, discarding the first 3,000 iterations of each chain as burn-in and using no thinning, which we considered appropriate for the complexity of our model (Gelman et al., 2013). To assess model fit and predictive ability, we conducted a two-stage model assessment. First, we assessed all models using fitted Bayesian R² to quantify the explained in-sample variance (Gelman et al., 2019). We also used the expected log pointwise predictive density (elpd) in Leave-One-Out cross-validation (LOO) as an approximation to out-of-sample predictive accuracy (Vehtari et al., 2024). In a second stage, we evaluated predictive performance using Leave-Future-Out cross-validation (LFO), which assesses out-of-sample predictive accuracy in a time-series context (Bürkner et al., 2020). Please refer to Supplementary Methods SM3.2 for details and references for acceptance thresholds and the model evaluation tools used.

### 5. Sensitivity analysis

We divided the sensitivity analysis into two parts. First, we applied eleven alternative prior specifications to the highest-scoring model to evaluate prior sensitivity (Table SM2). Each variation involved either a left shift (i.e., decrease) in a vital rate or a change in the shape of its prior distribution. Narrowing reflects increased prior confidence (reduced uncertainty), whereas flattening represents reduced prior confidence. We focused only on left shifts, as most posteriors are strongly right-skewed; a right shift would likely generate biologically implausible results. We grouped the variations into two categories: global variations that modify all vital rates or haul-out parameters simultaneously, and local variations that target specific processes.

Secondly, we tested the sensitivity of the results to alternative functional forms by re-fitting the highest-scoring model using three additional formulations. The original model used a cloglog link with quadratic covariate terms. We compared this to three variants: (1) cloglog with a linear term, (2) logistic with a linear term, and (3) logistic with quadratic terms. This comparison allows us to evaluate how the choice of link function influences model fit and inference.

### 6. Projections

#### 6.1. ENSO scenarios

To assess the extirpation risk of *O. byronia* in Peru under anticipated increases in extreme El Niño events, we projected population size over 50 years (2023–2072) using the full posterior from the fitted state-space model under six scenarios. Under the high-emission scenario SSP5-8.5 (IPCC, 2021), the frequency of extreme El Niño events is projected to rise by 59%, from 5.57 ± 0.81 events during 1900–1999 to 8.81 ± 1.03 in 2000–2099 (Cai et al., 2021), with approximately 25% intensification of ENSO variability already observed between the early and late twentieth century (Cai et al., 2021; Huang et al., 2017; Ishii et al., 2005; Rayner et al., 2003). The frequency of extreme La Niña events is also projected to rise by 48%. Precisely translating these long-term projections into discrete changes in event frequency and intensity remains challenging. We therefore defined six scenarios isolating and combining the effects of frequency and intensity across three levels each, using the observed ENSO regime during the model fitting period (1997–2022) as the baseline (Table 1). We generated stochastic covariate time series under each of the six defined ENSO scenarios, with the covariate used in the highest-scoring model (details in Supplementary Methods SM4.1). These independent time series were then used as inputs to a forward projection model with the same structure as the process model. To propagate parameter uncertainty through the projections, we ran the model using all posterior samples of the population size in 2022 and baseline vital rates. To propagate environmental stochasticity, we generated annual environmental errors and assigned a unique covariate time series to each sample. This produced 11,998 simulations per scenario over a 50-year projection period, incorporating uncertainty in model parameters, environmental stochasticity, and variability in climatic projection.

**Table 1.** ENSO projection scenarios used in population simulations.

| Scenario | El Niño<br>frequency<br>change | La Niña<br>frequency<br>change | Intensity<br>change | Projection basis |
| --- | --- | --- | --- | --- |
| Baseline | None | None | None | Reflects observed ENSO regime during model fitting period; assumes projected intensification for this century has already occurred and will persist |
| Medium frequency | +30% | +24% | None | Approximates half of the projected end-of-century increase in event frequency <sup>a</sup> |
| High frequency | +59% | +48% | None | Reflects full end-of-century increase by 2072; upper-bound scenario for sensitivity analysis <sup>a</sup> |
| Medium intensity | None | None | +10% | Approximates half of the intensification observed between the early and late 20th century <sup>b</sup> |
| High intensity | None | None | +20% | Upper-bound intensity scenario <sup>b</sup> |
| Combined | +30% | +24% | +10% | Moderate change in both dimensions; represents a plausible mid-range future condition |

<sup>a</sup> (Cai et al., 2021)
<sup>b</sup> (Cai et al., 2021; Huang et al., 2017; Ishii et al., 2005; Rayner et al., 2003)
Note: Scenarios do not imply a linear progression of ENSO change through time, but rather bracket plausible future conditions given uncertainty in the timing and trajectory of projected increases.

**Table 2.** Mean estimated extirpation risk and associated 95% Wilson score confidence intervals (derived from a single set of posterior projection simulations) for the Peruvian *O. byronia* population under six ENSO scenarios, assessed as the probability of meeting the IUCN Red List A2 criterion threshold for Critically Endangered during the projection period.

|  |  | Lower | Upper |
| --- | --- | --- | --- |
|  | Mean | 95% CI | 95% CI |
| Original | **0.25** | 0.24 | 0.26 |
| Medium frequency | **0.34** | 0.33 | 0.35 |
| High frequency | **0.41** | 0.40 | 0.42 |
| Medium intensity | **0.35** | 0.34 | 0.35 |
| High intensity | **0.44** | 0.43 | 0.45 |
| Combined (Medium frequency and intensity) | **0.46** | 0.45 | 0.47 |

#### 6.2. Extirpation risk metrics

To provide a more comprehensive assessment of extirpation risk, we applied the IUCN Red List Criterion A2 (IUCN, 2012) as a population-level benchmark. We estimated the probability that the Peruvian population meets the Critically Endangered threshold, defined as a ≥80% decline over three generations. Another threshold option is effective population size (Ne), for which de Oliveira et al. (2012) provide a demographically derived estimate for the Peruvian population. However, this estimate is not based on genetic data, and adapting it to a female-only threshold would require additional assumptions that introduce uncertainty. We therefore rely on Criterion A2 as our sole threshold, see Supplementary Methods SM4.2 for details.

## Results

Model selection identified an SST-based model as the best descriptor and predictor of the historical population dynamics of *O. byronia* in Peru. We present the model selection process and reconstructed population time series, followed by estimated vital rates and the relative contributions of environmental drivers, density dependence, and stochasticity to their variation. We then present population projections under six ENSO scenarios evaluated against effective population size thresholds and the IUCN A2 criterion, including a sensitivity analysis of projection outcomes. Finally, we compare two correction multipliers used to account for unobserved individuals in survey counts.

### 1. Model selection and population reconstruction

#### 1.1. Best model and reconstructed time series

We identified the best SST-based model using a two-stage approach integrating three complementary performance metrics. In the first stage, both LOO and fitted R² indicated that all ten top-performing models used covariates from the 12 months preceding the breeding season for survival rates. No single model was statistically distinguishable from the others within this group, and LOO estimates showed possible bias with 4.2–37.5% of Pareto k values exceeding 0.7 (see Discussion). We therefore proceeded to the second stage with these ten models and the base model. LFO cross-validation, which is more appropriate for time-series evaluation as it tests predictive ability on future observations (Bürkner et al., 2020), identified an SST-based model (model 19) as the top performer, marginally outperforming the leading zooplankton model (elpd difference: −1.50 ± 1.20). This SST model applied covariates from the 12 months preceding the breeding season for both survival rates and birth rate. Although standard errors were wide, it ranked joint first in stage one. We therefore selected this model for subsequent projections (see Supplementary Results SR1 for full model selection details).

The reconstructed time series with the selected SST-based model effectively captures the sharp population declines associated with the two strongest El Niño events during the 26-year study period: 1997–1998 and 2015–2016 (Figure 4). The estimated total female population was 205,457 (90% CrIs: 154,920 - 289,055) in 1997. It dropped sharply to 60,972 (44,640 - 82,678) in 1998, representing a 70.3% decline. The population then gradually increased, reaching a peak of 175,036 (130,230 - 234,380) in 2014. This corresponds to a compound annual growth rate of approximately 6.8% between 1998 and 2014, with a highest observed annual growth rate of 15.0% in 2000. Following the peak in 2014, the population declined again to 129,116 (98,514 - 169,695) in 2016, a 26.2% decrease from 2015. The SST model captures the population peak before the 2015–2016 El Niño event, despite a data gap in the years prior. The SST model also shows substantially smaller uncertainty in its estimates than the base model, as the latter relies solely on an environmental stochasticity term to account for interannual variability.

**Figure 4.**
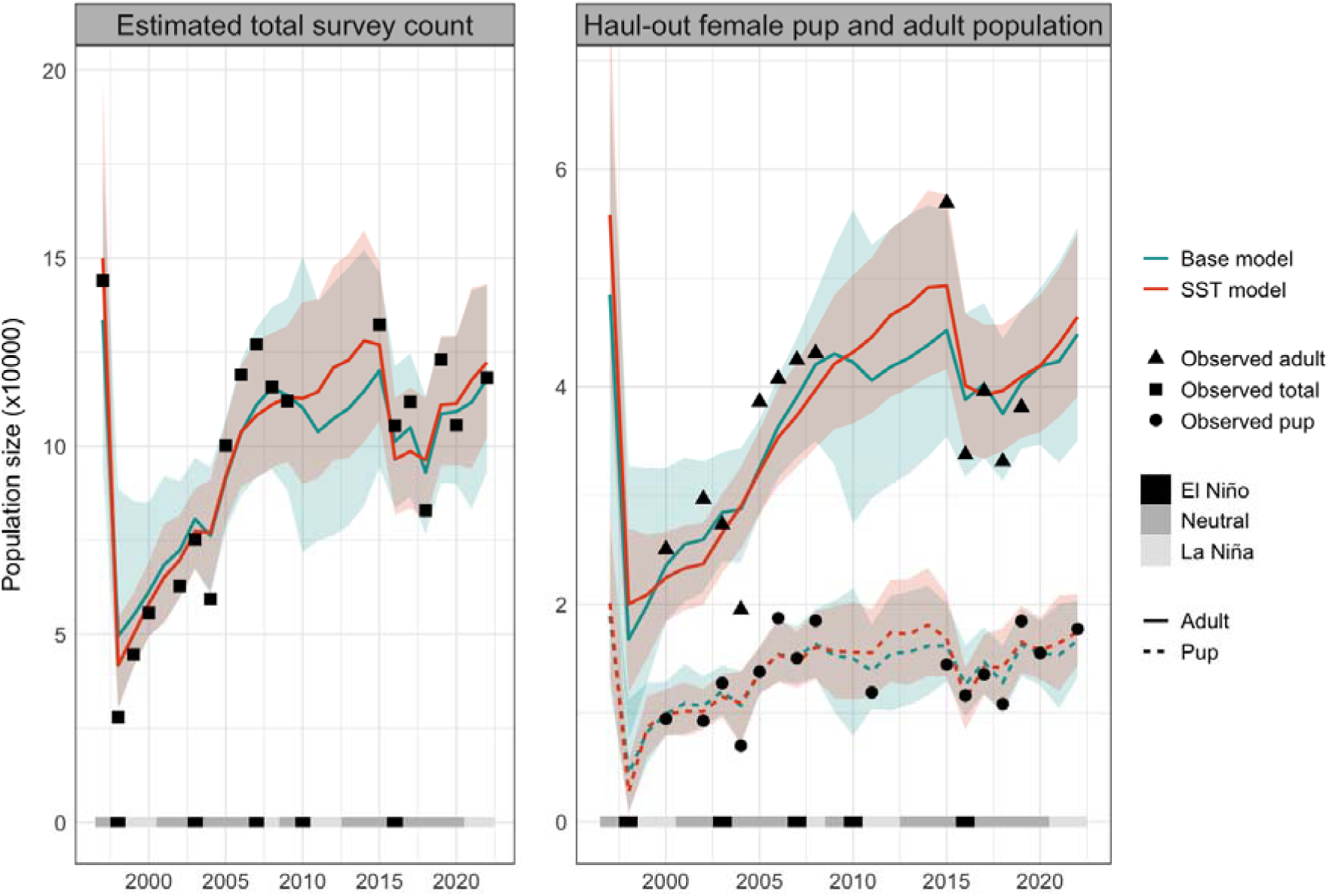
Reconstructed time series of population size and observed counts. The left panel shows the model-estimated total population count (both sexes) and the right panel displays haul-out counts derived from the stage-specific haul-out ratio as estimated by the model. In the right panel, the dashed line represents the female pup population that hauls out and the solid line represents the female adult population that hauls out. The coloured shade around the line represents the 90% CrIs for the estimated number. The two colours correspond to the different models. The black points indicate raw data points, and the grayscale -, and neutral years, with classifications sourced from NOAA’s Physical Sciences Laboratory (https://psl.noaa.gov/enso/past_events.html).

#### 1.2. Vital rate estimates

We present the first estimates of key vital rates and haul-out ratios for this population, providing a reference for future studies. Posterior estimates for survival and birth rates mostly fall on the higher end of their prior distributions, except for pup survival (Figure SR3). This pattern is driven by the assumed negative density dependence and the strong negative covariate effects incorporated through the complementary log-log link, both of which tend to suppress yearly vital rates. To better approximate vital rates under moderate population sizes and neutral environmental conditions, we report the average estimated vital rates across years in Table 3. The average yearly pup survival and juvenile survival are both lower than the prior mean. Posterior estimates of the mean haul-out ratios also differ considerably from their priors: the pup haul-out ratio is 21.3% lower, while the adult female ratio is 50.0% higher than the prior mean.

**Table 3.**
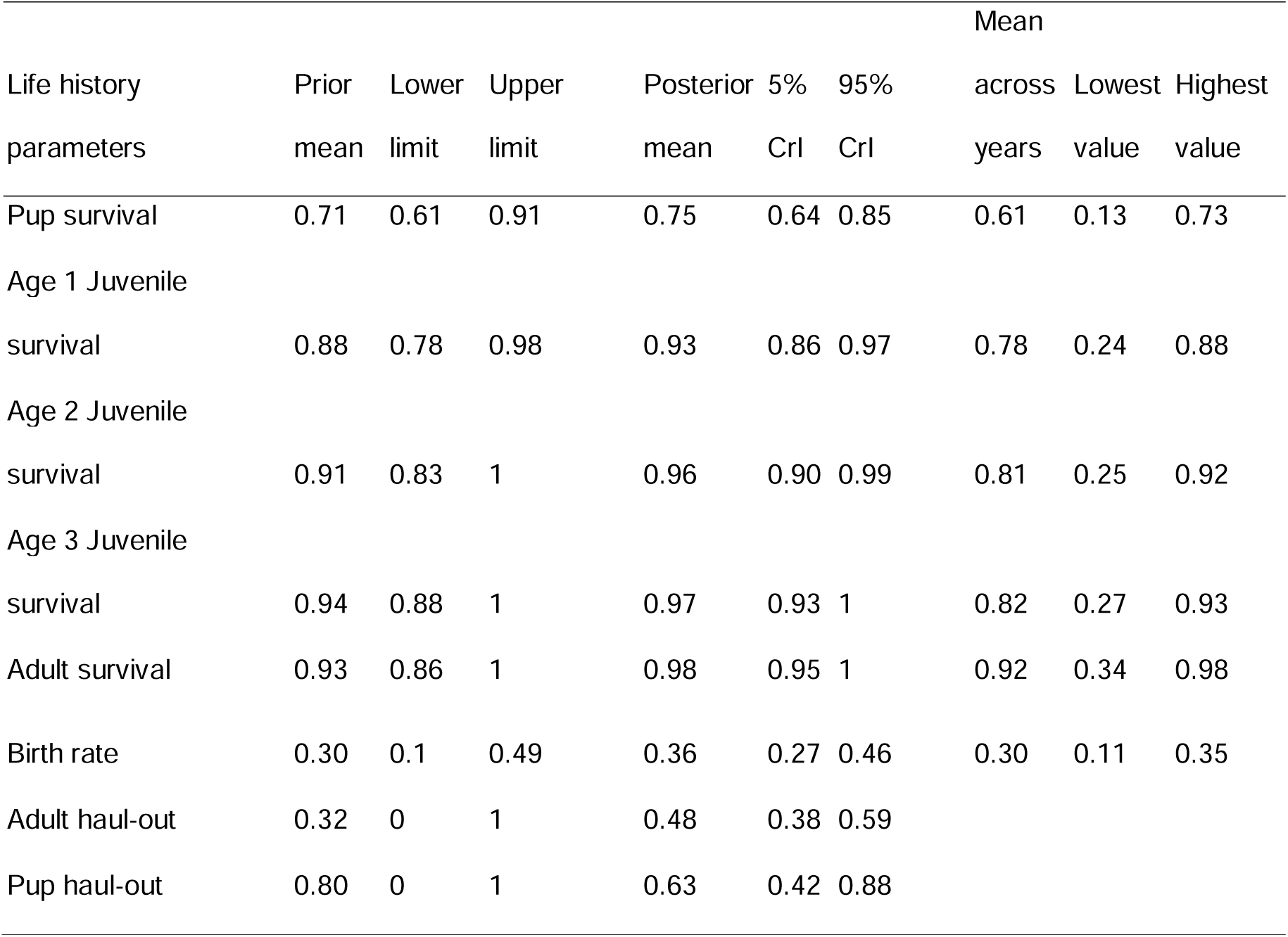
Prior distributions, posterior distributions, and yearly estimate summaries for *O. byronia* vital rates and haul-out ratios. The ranges represent different concepts: the prior range shows the lower and upper bounds of the prior distribution, the posterior range reflects the 90% credible intervals (Crl), and the yearly range indicates the minimum and maximum of yearly posterior mean estimates. Posterior mean estimates are higher than prior means because they represent the base rate, which is subsequently reduced by negative density-dependent regulation and covariate effects. Note that the birth rate represents female pups only.

#### 1.3. Drivers of vital rate variation

To assess how covariate, density dependence, and environmental stochasticity influence population dynamics, we modelled their effects on vital rates. Covariate effects strongly influenced all vital rates during the two major El Niño years (1997–1998 and 2015–2016; Figure 5). The absolute decrease in birth rates during the two major El Niño years appears smaller than the decrease observed in other demographic rates because baseline birth rates are lower, reflecting female-only reproduction. However, the proportional reduction is substantial, and actual births are further reduced by high adult mortality during these periods, which lowers the number of breeding females. Covariate effects account less well for smaller interannual fluctuations, particularly in pup survival. Environmental stochasticity, representing residual interannual variation not captured by the measured covariates, exerts a strong influence on pup survival but has a minimal impact on adult survival. In years such as 2004, unexplained variation in pup survival was largely attributable to this residual stochastic component. Density dependence has the least effect on adult survival but exerts a moderately consistent influence across the other vital rates, contributing to a gentle downward trend in the vital rates when the population increases.

**Figure 5.**
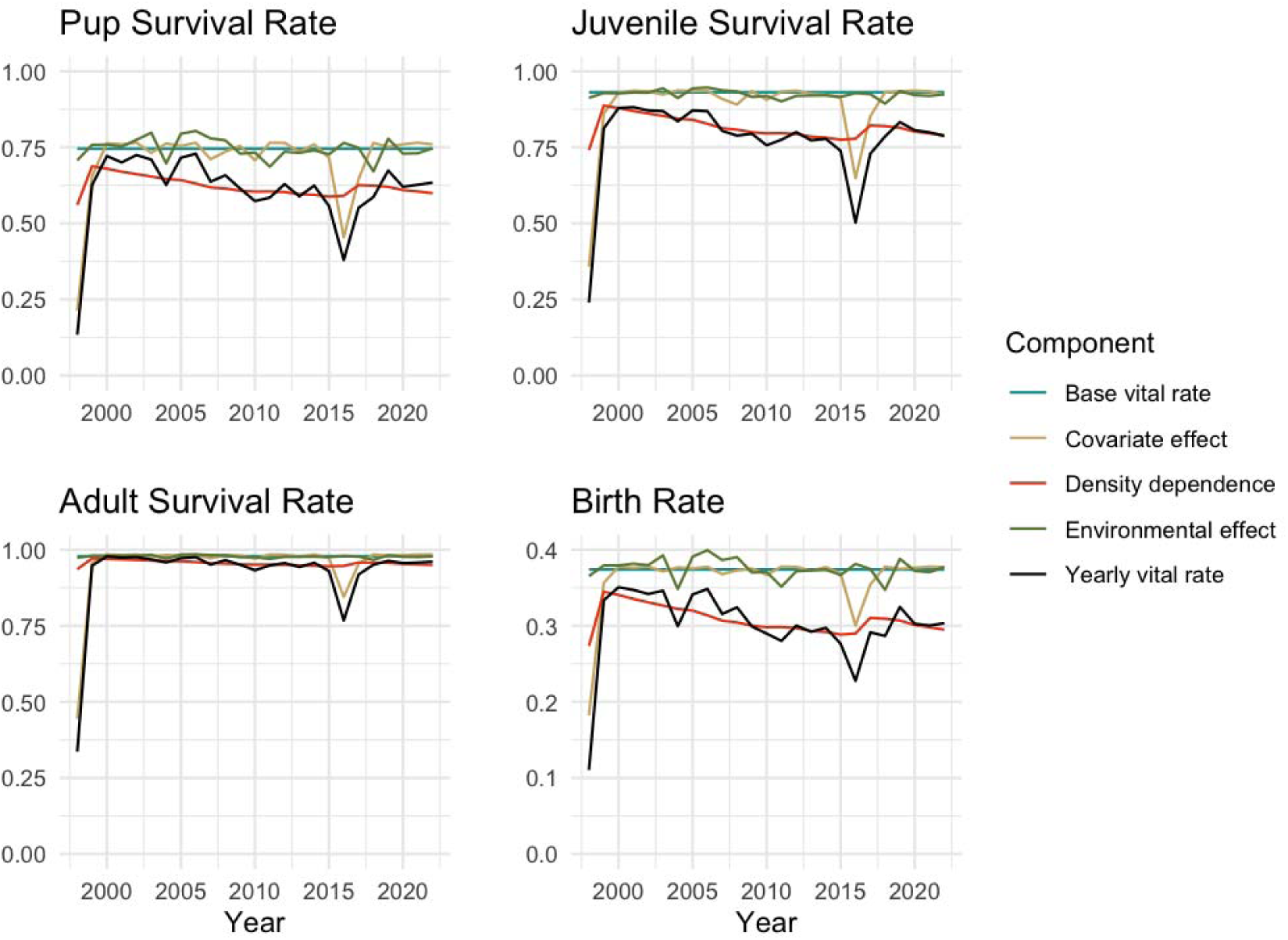
Visualization of the yearly and isolated effects of covariates, environmental stochasticity, and density dependence on the birth rate and the survival rates of pups, age-1 juveniles, and adults. The black line shows the posterior yearly mean for each vital rate, extracted directly from the SST model. Isolated effects are computed using posterior means of model parameters and differ from the posterior mean by less than 0.001% in all cases. Note that the y-axis scales differ between panels. Only the survival rates of age-1 juveniles are shown; survival patterns for age-2 and age-3 individuals are similar.

### 2. Population Projections and Extirpation Risk

#### 2.1. Projected population dynamics

Despite historical recovery under past ENSO regimes, projected alterations to ENSO dynamics substantially increase the extirpation risk of the Peruvian *O. byronia* population. Over the 50-year projection horizon, every altered ENSO scenario results in a higher extirpation probability than the baseline (original) scenario under the IUCN criterion. Greater event intensity consistently increased the risk of extirpation more than greater event frequency (Table 2). For example, the High-Intensity scenario increased the extirpation risk by 73.9%, whereas the High-Frequency scenario increased it by 62.7%. The Combined-Medium scenario produced the highest overall risk, surpassing all single-factor scenarios; the extirpation probability was 83.4% higher than the baseline.

#### 2.2. Sensitivity analysis

Extirpation risk projections were robust to alternative prior specifications but were sensitive to assumptions about the functional form used to model covariate effects. Although quantitative estimates varied across model structures, the qualitative pattern was consistent: extirpation risk increased under intensified El Niño conditions in all cases. Results for model-structure sensitivity are presented below (more details in Supplementary Results SR3), while prior sensitivity analyses are provided in SR2.

Alternative functional forms led to a marked divergence in extirpation risk metrics. The original formulation, using a complementary log-log link with quadratic covariate terms, achieved the highest elpd in both LFO and LOO comparisons, with overlapping confidence intervals. In contrast, the logistic link with a linear covariate term resulted in the poorest fit to the data. Examination of yearly mean vital rates revealed that the logit–linear formulation estimated higher juvenile survival, which buffered total abundance against steep declines and substantially reduced the absolute baseline extirpation risk by half. However, all climatic scenarios increase extirpation risk relative to the baseline, with the combined scenario showing a 94.6% higher risk than the baseline scenario.

### 3. Correction multipliers

The total count multiplier, calculated as the ratio of estimated total female (adult and pup) population to observed total counts (both sexes), appeared to be a better proxy for estimating true female abundance compared to the pup-based multiplier. Using GAMs that model the relationship between each multiplier and estimated female population, the total count multiplier exhibited a significant non-linear relationship with estimated female abundance (edf = 1.83, F = 7.05, p = 0.01); this model explained 45.3% of the deviance (adjusted R² = 0.39). In contrast, the pup-based multiplier showed no significant association (edf = 1, F = 0.03, p = 0.87), with negligible explanatory power (adjusted R² = –0.07). Furthermore, the total count multiplier demonstrated substantially lower variance (range: 1.12–2.18, mean: 1.42, SD: 0.26) than the pup-based multiplier (range: 7.34-15.78, mean: 10.17, SD: 2.25), indicating greater stability. These results suggest that, for highly fluctuating populations like *O. byronia* in Peru, total observed counts may serve as a more reliable proxy for correcting observed abundances to the estimated true population size than pup counts. However, we suggest caution, as this finding may be impacted by the model’s lower ability to capture fluctuations in pup abundance.

## Discussion

We assessed how increased El Niño frequency will impact the projected extirpation risk of *O. byronia* in Peru over the next 50 years. To generate this estimate, we constructed a state-space population model to evaluate how environmental variability has shaped population dynamics in Peru over the past 26 years. We also quantified shifts in vital rates during two major El Niño events and identified SST anomalies as the best predictor of historical population trajectories. Under the high-emission scenario SSP5-8.5, extreme El Niño events in this century are expected to occur 60% more frequently than in the 20th century (Cai et al., 2021). Our most conservative frequency scenario assumes that half of the projected end-of-century increase has already occurred, corresponding to a remaining 30% increase in frequency. Even under this mild assumption, our projections indicate a 36.1% increase in the risk of the population crossing the extirpation threshold. When this is combined with a modest 10% increase in El Niño intensity, an observed trend over recent decades (Lee & McPhaden, 2010), the risk rises by 83.4%. Beyond the frequency and intensity of ENSO events, climate-driven changes in prey dynamics pose an additional threat. Bas et al. (2025) projected that, for the Pacific population of *O. byronia*, climate-driven changes will reduce spatial overlap with key prey species and lead to a shift toward lower-quality prey resources. Although the population has shown apparent resilience in the past two decades, continued warming and increasingly extreme oceanographic conditions could ultimately exceed the species’ adaptive capacity.

In addition to climate change, emerging disease threats have raised new concerns for the Peruvian *O. byronia* population. The first recorded outbreak of highly pathogenic avian influenza (HPAI) A(H5N1) in *O. byronia* occurred in Peru between January and March 2023, during which 11,000 dead individuals were observed (Kuiken et al., 2026). The virus is believed to have been introduced by migratory birds traveling from North America to Central and South America, during a broader panzootic that affected wildlife globally from 2020 to 2023 (Gadzhiev et al., 2024; Wille & Barr, 2022). Infectious disease-induced mass mortality events have been positively correlated with global annual SST anomalies, suggesting an elevated risk of future outbreaks under continued climate warming (Sanderson & Alexander, 2020). Other human-related mortality sources have also been documented. A mass mortality event in 2009 in Piura was linked to pesticide exposure, likely due to intentional poisoning (IMARPE, 2009). High concentrations of contaminants such as aluminum, arsenic, and lead have been reported in this population, suggesting potential effects on immune and reproductive health (Kooyomjian et al., 2022). The operational interactions between *O. byronia* and fisheries are substantial, with over 4,800 individuals recorded interacting with the Peruvian industrial purse-seine fleet between 2017 and 2022 (Peraltilla et al., 2024), posing elevated bycatch risk to sea lion populations (Sepúlveda et al., 2023; Sepúlveda et al., 2026). Higher likelihoods of harmful encounters with fisheries were observed during El Niño years, as indicated by higher stranding rates of *O. byronia* showing signs of blunt trauma and poisoning (Bachmann Caller, 2018). Climate intensification may amplify these existing threats, creating compound pressures that exceed the population’s buffering capacity.

Vital rates estimated from the best SST model reasonably capture population dynamics under normal conditions and robustly quantify the severe impact of extreme climatic events on *O. byronia*. The estimated adult survival during the 1997–1998 El Niño event was 0.34, approximately aligning with field-based estimates (∼0.4). However, the model’s pup and juvenile survival estimates for the same year (0.13–0.27) are higher than field-based estimates in these age classes (Soto et al., 2004), though that study was limited to a single rookery and may not reflect population-level survival. Importantly, the highest and mean annual estimates of pup survival, juvenile survival, and reproduction all fall within the range observed for other otariid species (Table SR8). Our lowest annual estimates are lower than all previously reported values. For instance, Melin et al. (2012) reported a total birth rate of 0.37 (SE = 0.11) for California sea lions during the 1997–1998 El Niño event, among the less productive age group (17–21 years). Assuming a 1:1 sex ratio, this translates to 0.19 female pups per female, which is still 72.7% higher than our lowest estimate of 0.11 (90% CrIs: 0.02–0.24) for the same year. These findings demonstrate the uniquely challenging position of *O. byronia* at the epicenter of ENSO variability.

A key factor in coping with this challenge is the high survival of adult females. Our highest annual stage-based estimate for adult survival reached 0.98, exceeding the upper range reported for most other otariid species. For comparison, adult female survival in a growing *O. byronia* population in Argentina was 0.93 (Grandi et al., 2016). Across all our sensitivity scenarios, except those where the prior on adult survival was explicitly lowered, the models consistently favoured high adult female survival. When we lowered the adult survival, other vital rates changed to compensate. This compensation came at the cost of biological plausibility – for example, the predicted pup haul-out rate dropped further, contradicting general expectations of high pup haul-out (Grandi et al., 2021). Model estimates show that adult survival in the Peruvian *O. byronia* was comparatively stable during moderate environmental fluctuations from 2002 to 2005, when pup survival and reproduction fluctuated, consistent with demographic buffering being more sensitive to environmental variance than autocorrelation (Gascoigne et al., 2025; Koons et al., 2009). This stability may be supported by the high foraging plasticity of lactating *O. byronia* females (Cárdenas-Alayza et al., 2022b) and the life-history strategy of early weaning to prioritize adult survival, which was anecdotally observed in this species (Trivers 1974; Sepúlveda & Harcourt, 2021). However, high estimated female survival could partly be confounded with increased immigration from Chile during favourable conditions, given the model assumption of a closed population.

We demonstrated that top predator abundance is strongly influenced by low-trophic-level conditions and rapid response dynamics, consistent with the generally bottom-up-driven nature of the Humboldt Current System. The best SST model used the 12-month pre-breeding window to explain variation in survival and birth rate, and had better predictive performance than zooplankton models, likely stemming from its stability and lower variability than zooplankton. Both SST and zooplankton were better predictors than anchovy, despite earlier literature emphasizing anchovy as a key prey species (Soto et al., 2006; Arias-Schreiber, 2003). While anchovy, red squat lobster, cephalopods, and various fish species are all major components of the diet of *O. byronia* (IMARPE 2005, 2006; Cárdenas-Alayza et al., 2022a), dietary composition varies markedly across regions and over time (IMARPE 2000, 2009). For example, during the 2015 El Niño event, red squat lobster consumption at Punta San Juan dropped from an average of 69.9% to just 2% (Cárdenas-Alayza et al., 2022a). This variability suggests that SST or zooplankton may serve as more consistent proxies for energy flow through the food web. However, this result should be interpreted with caution, as the anchovy data were only available at a coarser temporal resolution (seasonal rather than monthly), which may reduce their accuracy in representing yearly biomass.

The estimated vital rates and covariate effects are biologically plausible, despite two major challenges in the model selection process. These challenges were, computationally, the absence of fast approximation methods tailored for time series data and, statistically, the high uncertainty in our evaluation metrics. During LOO analysis, all models produced Pareto k values exceeding 0.7, a known issue where individual observations exert disproportionate influence on parameter estimates (Vehtari et al., 2017), potentially underestimating predictive accuracy. Such values can arise when models are highly flexible or when priors are weakly informative, complicating the prediction of left-out observations even if the model is correctly specified (Vehtari et al., 2024). In our case, better-fitting models with higher LOO scores, which captured rapid fluctuations in the time series more closely, also tended to have higher Pareto k values due to these influential observations. To address this, we conducted exact LFO cross-validation, accepting the increased computational cost given the absence of an off-the-shelf implementation. A further obstacle was the wide credible intervals of all three evaluation metrics (LOO, LFO, and Bayesian R²), reflecting the sparse, noisy nature of the observational data, the complexity of the model, and the use of wide priors for vital rates. Nevertheless, posterior predictive checks confirmed good agreement between observed and predicted counts for the best SST model. We acknowledge that this uncertainty is inherent to modelling ecological dynamics with limited, noisy data.

Accurate population estimates are essential for assessing population risk and can be obtained through calibrated extrapolation from counts. We found a significant relationship between the total count multiplier and female abundance, but not for the pup-based multiplier. Our mean estimate of the total count multiplier across years is 1.42 (range: 1.12–2.18). Because this estimate uses total counts (both sexes) to infer the true female population, the corresponding multiplier for the total population would be higher. This aligns with findings from other pinniped populations, where reported multipliers are larger, such as 1.75–1.89 for *O. byronia* in Argentina (Romero et al., 2017). Our mean estimate of the pup-based multiplier across years is 10.17 (range: 7.34–15.78). This value is higher than those reported for other pinnipeds (3.00–7.71 across 19 populations; Karamanlidis, 2024), likely reflecting the very low pup production observed during El Niño years. We suggest that extrapolating total population counts (both sexes) provides a more reliable estimate of the true female population, offering greater stability over time. This is particularly true when accounting for the inverse, non-linear relationship between the multiplier and population size. This is likely to hold for other highly fluctuating populations, where pups are the most affected group during poor years, and adults are impacted only in extreme years.

An alternative approach to estimating pinniped population sizes relies on estimating haul-out ratios, but uncertainty remains a major source of error. To capture variation in pup presence, we defined the haul-out ratio on any given date in the breeding season based on weekly data from a single site. We also assumed that all pups are detected during the peak of the breeding season and pupping phenology is constant across the year, which is unlikely to be true (Soto et al., 2004). The variance of the resulting prior is therefore higher than expected. This issue could be improved with weekly or even daily counts across all colonies, though such data collection is highly challenging in the context of a national census. For adults, estimating the haul-out ratio is complicated by their variable haul-out behavior throughout the day (Sepúlveda et al., 2012). In our case, this may be less problematic, as censuses are consistently conducted in the morning. Variation across years, however, is more difficult to address. For simplicity, we assumed a constant adult haul-out ratio across years, even though prey availability can influence foraging trip duration and consequently reduce the haul-out ratio (Soto et al., 2004). In general, improving model performance will require more informative priors with greater confidence (i.e., lower variance), which can be achieved by incorporating better empirical data across multiple parameters.

Our projections highlight a broader challenge for conservation under accelerating climate change. Management frameworks based on static abundance thresholds assume environmental stationarity, yet our results demonstrate that intensifying disturbance regimes can greatly shift extirpation risk even in historically resilient populations. In such systems, reference points such as the Maximum Net Productivity Level (MNPL) used in US marine mammal management, which are often treated as static, may offer limited guidance. In Peru, the stated objective of the marine reserve system (RNIPG) is to ‘conserve a representative sample of the marine biodiversity from the Northern Humboldt Current Upwelling System’ (Del Solar et al., 2023), yet what constitutes a “representative sample” for *O. byronia* remains undefined. Our projections point to the need for precautionary management that explicitly accounts for the risk of episodic collapse. In particular, we propose two broad management priorities, which are monitoring and forage protection.

First, maintaining long-term monitoring efforts and improving transparency in data reporting can enhance early detection of demographic change and reduce uncertainty in population assessments. This approach includes publishing detailed survey metadata, e.g., dates and times, disaggregated regional and life-stage data, and explicit measures of uncertainty, including confidence intervals. Additional individual-based studies, including satellite tagging and mark-recapture experiments, could provide more informative priors. Second, strengthening foraging protection by establishing true no-take zones in key breeding habitats could buffer populations against repeated disturbances (Gormley et al., 2012). In Peru, for example, illegal activities such as dynamite fishing have been reported near the Paracas National Reserve (IMARPE, 2004; 2007); within the RNIPG, only a two-nautical-mile boundary is protected from fishing, leaving adjacent habitats vulnerable (Laínez del Pozo & Jones, 2021). This strategy should also be supported by transparent, precautionary, and ecosystem-based industrial fisheries quotas, with a focus on reducing trophic competition between predators and fisheries during extreme climatic events. Together, these measures support the objective of conserving a truly representative and resilient *O. byronia* population.

These directions, monitoring and forage protection, are also broadly relevant to marine predators exposed to intensifying climate extremes. The international conservation movement of the 1970s, marked by landmark legislation such as the Marine Mammal Protection Act (1972) and CITES (1973), demonstrated that coordinated policy could reverse population declines by addressing a primary, identifiable pressure – direct harvesting (Magera et al., 2013). Climate change, however, operates at a fundamentally different scale, acting through multiple, interacting pathways that no single legislative act can eliminate. As disturbance regimes intensify globally, resilience-oriented management frameworks grounded in species-specific demographic data and environmental context, as demonstrated here, will be essential to safeguard marine mammal populations against an increasingly volatile climate.

## Supporting information

Supplementary Results 3

Supplementary Results 4

## Author Contributions

HWYC conceived the original idea, which was further developed with critical input from KJD and MBB. HWYC conducted the analysis and led the writing of the manuscript. SC, MJG, and KFH provided data from Punta San Juan, which were essential for prior specification, and contributed to the review of methods and results. JA and JCM facilitated in requesting part of the count data through the transparency platform from Instituto del Mar del Perú (IMARPE). All authors contributed critically to the revisions and approved the final manuscript for publication.

## Acknowledgements

We would like to express our gratitude to the information received from IMARPE (N° 149-2020 and 070-2020-IMARPE/OGA), through the Ley de Transparencia y Acceso a la Informacion Publica Ley 27806 and DS 043-2003 PCM, and to the staff responsible for the long-term country-wide surveys. We would like to thank the consortium of zoological organizations that make long-term data collection by the Punta San Juan Program. We also thank Dr. Tim Tinker for his help in transitioning our original JAGS script to Stan.

## Data Availability Statement

Code, Stan model files, supplementary workflows, processed covariate inputs, and archived model outputs supporting this study are publicly available on Zenodo at [https://doi.org/10.5281/zenodo.20394833]. Raw count and telemetry data from Punta San Juan used to inform prior specification are the property of collaborating institutions and cannot be publicly archived; these data are available from the corresponding author upon reasonable request and with permission from the data providers. All data necessary to reproduce the findings of this study are contained within the Zenodo deposit.

## ## List of Supplementary Materials

### ### Supplementary Methods, Data & Results 1–2

Word document containing detailed descriptions of:

SM1. Model structure and assumptions

SM2. Covariate sources and processing

SM3. Bayesian model implementation and evaluation

SM4. Projection framework

SD1. Compilation and harmonisation of count data from multiple sources

SR1. Model selection

SR2. Prior sensitivity analysis

### ### Supplementary Results 3

Excel document containing:

- Table SR1. Model results

- Table SR2. Fitted Bayesian R²

- Table SR3. Leave-future-out (LFO) expected log predictive density (ELPD) results

- Table SR4. Prior sensitivity analysis

- Table SR5. Population trajectory summaries

- Table SR6. Functional form sensitivity analysis

- Table SR7. Projection sensitivity analysis

### ### Supplementary Results 4

Excel document containing:

- Table SR8. Reported ranges of female vital rates across otariid species

## References

Aarts, G., Brasseur, S., Poos, J. J., Schop, J., Kirkwood, R., Van Kooten, T., Mul, E., Reijnders, P., Rijnsdorp, A. D., & Tulp, I. (2019). Top-down pressure on a coastal ecosystem by harbor seals. Ecosphere, 10(1), e02538. 10.1002/ecs2.2538

Alam, E., & Linero, A. R. (2025). A Unified Bayesian Nonparametric Framework for Ordinal, Survival, and Density Regression Using the Complementary Log-Log Link. arXiv preprint arXiv:2502.00606. 10.48550/arXiv.2502.00606

Arias-Schreiber, M., & Rivas, C. (1998). Distribución, tamaño y estructura de las poblaciones de lobos marinos arctocephalus australis y otaria byronia en el litoral peruano, en noviembre 1996 y marzo 1997 (No. 73). Instituto del Mar del Perú.

Arias-Schreiber M (2003) Prey spectrum and feeding behaviour of two sympatric pinnipeds (Arctocephalus australis and Otaria flavescens) in relation to the 1997–98 ENSO in southern Peru. MSc thesis, University of Bremen

Avila, I. C., Kaschner, K., & Dormann, C. F. (2018). Current global risks to marine mammals: taking stock of the threats. Biological Conservation, 221, 44–58. 10.1016/j.biocon.2018.02.021

Bachmann Caller, V. M. (2018). Analysis of sea turtle and marine mammal strandings along the northern coast of Peru between 2003 and 2016: Patterns and underlying causes (Master’s thesis, Universidad Andrés Bello). https://repositorio.imarpe.gob.pe/server/api/core/bitstreams/db65874d-3625-4034-8d93-0812b46908b1/content

Bakun, A., & Weeks, S. J. (2008). The marine ecosystem off Peru: What are the secrets of its fishery productivity and what might its future hold? Progress in Oceanography, 79(2–4), 290–299. 10.1016/j.pocean.2008.10.027

Bas, M., Ouled-Cheikh, J., Fuster-Alonso, A., Julià, L., March, D., Ramírez, F., Cardona, L., & Coll, M. (2025). Potential Spatial Mismatches Between Marine Predators and Their Prey in the Southern Hemisphere in Response to Climate Change. Global Change Biology, 31(2), e70080. 10.1111/gcb.70080

Bauer, H., Chapron, G., Nowell, K., Henschel, P., Funston, P., Hunter, L. T., … & Packer, C. (2015). Lion (Panthera leo) populations are declining rapidly across Africa, except in intensively managed areas. Proceedings of the National Academy of Sciences, 112(48), 14894–14899. 10.1073/pnas.1500664112

Berkson, J. M., & DeMaster, D. P. (1985). Use of Pup Counts in Indexing Population Changes in Pinnipeds.

Bürkner, P. C., Gabry, J., & Vehtari, A. (2020). Approximate leave-future-out cross-validation for Bayesian time series models. Journal of Statistical Computation and Simulation, 90(14), 2499–2523.

Cai, W., Santoso, A., Collins, M., Dewitte, B., Karamperidou, C., Kug, J.-S., Lengaigne, M., McPhaden, M. J., Stuecker, M. F., Taschetto, A. S., Timmermann, A., Wu, L., Yeh, S.-W., Wang, G., Ng, B., Jia, F., Yang, Y., Ying, J., Zheng, X.-T., … Zhong, W. (2021). Changing El Niño–Southern Oscillation in a warming climate. Nature Reviews Earth & Environment, 2(9), 628–644. 10.1038/s43017-021-00199-z

Caillat, M., Cordes, L., Thompson, P., Matthiopoulos, J., & Smout, S. (2019). Use of state- space modelling to identify ecological covariates associated with trends in pinniped demography. Aquatic Conservation: Marine and Freshwater Ecosystems, 29, 101–118. 10.1002/aqc.3130

Cappozzo, H. L., & Perrin, W. F. (2009). South American sea lion: Otaria flavescens. In Encyclopedia of marine mammals (pp. 1076–1079). Academic Press. 10.1016/B978-0-12-373553-9.00244-3

Caswell, H. (2001). Matrix population models: construction, analysis, and interpretation. In Matrix population models: construction, analysis, and interpretation (pp. 722–722).

Campagna C., & Boeuf B. J. L. (1988). Reproductive Behaviour of Southern Sea Lions. 10.1163/156853988X00539

Cárdenas-Alayza, S., Crespo, E., & Oliveira, L. (2016). Otaria byronia. The IUCN Red List of Threatened Species 2016: e.T41665A61948292. 10.2305/IUCN.UK.2016-1.RLTS.T41665A61948292.en

Cárdenas-Alayza, S., Adkesson, M. J., Edwards, M. R., Hirons, A. C., Gutiérrez, D., Tremblay, Y., & Franco-Trecu, V. (2022a). Sympatric otariids increase trophic segregation in response to warming ocean conditions in Peruvian Humboldt Current System. PLOS ONE, 17(8), e0272348. 10.1371/journal.pone.0272348

Cárdenas-Alayza, S., Adkesson, M. J., Gutiérrez, D., Demarcq, H., & Tremblay, Y. (2022b). Strategies for segregation during foraging in sympatric otariids of the Peruvian upwelling Humboldt Current System. Marine Ecology Progress Series, 702, 153–170. 10.3354/meps14203

Cárdenas-Alayza, S., Gutiérrez, D., & Tremblay, Y. (2021). Trends in sympatric otariid populations suggest resource limitations in the Peruvian Humboldt Current System. Marine Environmental Research, 169, 105349. 10.1016/j.marenvres.2021.105349

Chavez, F. P., & Messié, M. (2009). A comparison of Eastern Boundary Upwelling Ecosystems. Progress in Oceanography, 83(1), 80–96. 10.1016/j.pocean.2009.07.032

Chung, H. W. Y., Cárdenas-Alayza, S., Alfaro Shigueto, J., Farfán-Hernandez, K., Ganoza-Gallardo, M. J., Mengel, J. C., Davis, K., & Bonsall, M. (2026). Supplementary Code and Workflows for “From Recovery to Risk: Intensifying ENSO Events Drive Extirpation Risk in a Marine Predator”. Zenodo. 10.5281/zenodo.20394833

Conn, P. B., Johnson, D. S., Hoef, J. M. V., Hooten, M. B., London, J. M., & Boveng, P. L. (2015). Using spatiotemporal statistical models to estimate animal abundance and infer ecological dynamics from survey counts. Ecological Monographs, 85(2), 235–252.

Davis, K. J. (2022). Managed culls mean extinction for a marine mammal population when combined with extreme climate impacts. Ecological Modelling, 473, 110122. 10.1016/j.ecolmodel.2022.110122

De Oliveira, L. R., Fraga, L. D., & Majluf, P. (2012). Effective population size for South American sea lions along the Peruvian coast: The survivors of the strongest El Niño event in history. Journal of the Marine Biological Association of the United Kingdom, 92(8), 1835–1841. 10.1017/S0025315411001871

Del Solar, A., Alfaro-Cordova, E., Mendo, T., Ortega-Cisneros, K., Arias Schreiber, M., Grillo-Núñez, J., & Alfaro-Shigueto, J. (2023). Peru case study: big challenges for small Islands: management and governance of “lobos de Afuera” Islands in the Peruvian upwelling ecosystem. In Challenges in Tropical Coastal Zone Management: Experiences and Lessons Learned (pp. 303–322). Cham: Springer International Publishing.

Gabry, J., et al. (2025). CmdStanR: R interface to CmdStan (Version 2.36.0) [Computer software]. https://mc-stan.org/cmdstanr/

Gadzhiev, A., Petherbridge, G., Sharshov, K., Sobolev, I., Alekseev, A., Gulyaeva, M., Litvinov, K., Boltunov, I., Teymurov, A., Zhigalin, A., Daudova, M., & Shestopalov, A. (2024). Pinnipeds and avian influenza: A global timeline and review of research on the impact of highly pathogenic avian influenza on pinniped populations with particular reference to the endangered Caspian seal (Pusa caspica). Frontiers in Cellular and Infection Microbiology, 14. 10.3389/fcimb.2024.1325977

Gascoigne, S. J., Kajin, M., Tuljapurkar, S., Santos, G. S., Compagnoni, A., Steiner, U. K., … & Salguero-Gómez, R. (2025). Structured demographic buffering: A framework to explore the environmental components and demographic mechanisms underlying demographic buffering. Ecology Letters, 28(2), e70066. 10.1111/j.1600-0706.2009.16399.x

Gelman, A., Carlin, J. B., Stern, H. S., Dunson, D. B., Vehtari, A., & Rubin, D. B. (2013). Bayesian data analysis (3rd ed.). Chapman and Hall/CRC.

Gelman, A., Goodrich, B., Gabry, J., & Vehtari, A. (2019). R-squared for Bayesian regression models. The American Statistician.

Germishuizen, M., Vichi, M., & Vermeulen, E. (2024). Population changes in a Southern Ocean krill predator point towards regional Antarctic sea ice declines. Scientific Reports, 14(1), 25820. 10.1038/s41598-024-74007-1

Gormley, A. M., Slooten, E., Dawson, S., Barker, R. J., Rayment, W., du Fresne, S., & Bräger, S. (2012). First evidence that marine protected areas can work for marine mammals. Journal of Applied Ecology, 49(2), 474–480. 10.1111/j.1365-2664.2012.02121.x

Grandi, M. F., Dans, S. L., García, N. A., & Crespo, E. A. (2010). Growth and age at sexual maturity of South American sea lions. Mammalian biology, 75(5), 427–436. 10.1016/j.mambio.2009.09.007

Grandi, M. F., Dans, S. L., & Crespo, E. A. (2016). Improvement in Survivorship: The Key for Population Recovery? Zoological Studies, 55, e9. 10.6620/ZS.2016.55-9

Grandi, M. F., García, N. A., & Crespo, E. A. (2021). Recomendaciones metodológicas para los censos de lobos marinos comunes, Otaria flavescens. Ecología Austral, 31(3), 532–545. 10.25260/EA.21.31.3.0.1417

Hansen, J., Ruedy, R., Sato, M., & Lo, K. (2010). Global surface temperature change. Reviews of geophysics, 48(4). 10.1029/2010RG000345

Hammond, P. S., Francis, T. B., Heinemann, D., Long, K. J., Moore, J. E., Punt, A. E., Reeves, R. R., Sepúlveda, M., Sigurðsson, G. M., Siple, M. C., Víkingsson, G., Wade, P. R., Williams, R., & Zerbini, A. N. (2021). Estimating the Abundance of Marine Mammal Populations. Frontiers in Marine Science, 8. https://www.frontiersin.org/articles/10.3389/fmars.2021.735770

Harrison, X. A., Blount, J. D., Inger, R., Norris, D. R., & Bearhop, S. (2011). Carry-over effects as cs of fitness differences in animals. Journal of Animal Ecology, 80(1), 4–18. 10.1111/j.1365-2656.2010.01740.x

Harvey, B. P., & Moore, P. J. (2017). Ocean warming and acidification prevent compensatory response in a predator to reduced prey quality. Marine Ecology Progress Series, 563, 111–122. 10.3354/meps11956

Huang, P. (2017). Time-varying response of ENSO-induced tropical Pacific rainfall to global warming in CMIP5 models. Part II: Intermodel uncertainty. Journal of Climate, 30(2), 595–608. 10.1175/JCLI-D-16-0373.1

Hunt Jr, G. L., Stabeno, P., Walters, G., Sinclair, E., Brodeur, R. D., Napp, J. M., & Bond, N. A. (2002). Climate change and control of the southeastern Bering Sea pelagic ecosystem. Deep Sea Research Part II: Topical Studies in Oceanography, 49(26), 5821–5853. 10.1016/S0967-0645(02)00321-1

Kuiken, T., Vanstreels, R. E. T., Banyard, A., Begeman, L., Breed, A. C., Dewar, M., … & Wille, M. (2026). Emergence, spread, and impact of high-pathogenicity avian influenza H5 in wild birds and mammals of South America and Antarctica. Conservation Biology, 40(1), e70052. 10.1111/cobi.70052

IPCC. (2021). Climate Change 2021: The Physical Science Basis. Contribution of Working Group I to the Sixth Assessment Report of the Intergovernmental Panel on Climate Change (V. Masson-Delmotte, P. Zhai, A. Pirani, S. L. Connors, C. Péan, S. Berger, N. Caud, Y. Chen, L. Goldfarb, M. I. Gomis, M. Huang, K. Leitzell, E. Lonnoy, J. B. R. Matthews, T. K. Maycock, T. Waterfield, O. Yelekçi, R. Yu, & B. Zhou, Eds.). Cambridge University Press. 10.1017/9781009157896

IMARPE. (1997–2022). Anuario Científico-Tecnológico IMARPE [Annual scientific-technical reports]. Callao, Peru: Instituto del Mar del Perú. Retrieved from [https://repositorio.imarpe.gob.pe/handle/20.500.12958/6]

Ishii, M., Shouji, A., Sugimoto, S., & Matsumoto, T. (2005). Objective analyses of sea-surface temperature and marine meteorological variables for the 20th century using ICOADS and the Kobe Collection. International Journal of Climatology, 25(7), 865–879. 10.1002/joc.1169

IUCN. (2012). IUCN Red List categories and criteria: Version 3.1 (2nd ed.). IUCN. https://portals.iucn.org/library/node/10315

Juárez-Ruiz, A., Pardo, M. A., Hernández-Montoya, J. C., Elorriaga-Verplancken, F. R., Milanés-Salinas, M. D. L. Á., Norris, T., … & Heckel, G. (2022). Guadalupe fur seal pup production predicted from annual variations of sea surface temperature in the southern California Current Ecosystem. ICES Journal of Marine Science, 79(5), 1637–1648. 10.1093/icesjms/fsac097

Karamanlidis, A. A. (2024). Using ‘pup multipliers’ to estimate demographic parameters of Mediterranean monk seals in the eastern Mediterranean Sea. Endangered Species Research, 53, 261–270. 10.3354/esr01301

Koons, D. N., Pavard, S., Baudisch, A., & Jessica E. Metcalf, C. (2009). Is life-history buffering or lability adaptive in stochastic environments?. Oikos, 118(7), 972–980. 10.1111/j.1600-0706.2009.16399.x

Kooyomjian, C., Giarikos, D., Adkesson, M., & Hirons, A. C. (2022). Evaluation of trace element concentrations in the serum and vibrissae of Peruvian pinnipeds (Arctocephalus australis and Otaria byronia). The Journal of Wildlife Diseases, 58(3), 608–620. 10.7589/jwd-d-21-00104

Kortsch, S., Primicerio, R., Fossheim, M., Dolgov, A. V., & Aschan, M. (2015). Climate change alters the structure of arctic marine food webs due to poleward shifts of boreal generalists. Proceedings of the Royal Society B: Biological Sciences, 282(1814), 20151546. 10.1098/rspb.2015.1546

Kovacs, K., Aguilar, A., Aurioles, D., Burkanov, V., Campagna, C., Gales, N., Gelatt, T., Goldsworthy, S., Goodman, S., Hofmeyr, G., Harkonen, T., Lowry, L., Lydersen, C., Schipper, J., Sipilä, T., Southwell, C., Stuart, S., Thompson, D., & Trillmich, F. (2012). Global threats to pinnipeds. Marine Mammal Science, 28, 414–436. 10.1111/j.1748-7692.2011.00479.x

Laake, J. L., Lowry, M. S., DeLong, R. L., Melin, S. R., & Carretta, J. V. (2018). Population growth and status of california sea lions. The Journal of Wildlife Management, 82(3), 583–595. 10.1002/jwmg.21405

Laidre, K. L., Stern, H., Kovacs, K. M., Lowry, L., Moore, S. E., Regehr, E. V., Ferguson, S. H., Wiig, Ø., Boveng, P., Angliss, R. P., Born, E. W., Litovka, D., Quakenbush, L., Lydersen, C., Vongraven, D., & Ugarte, F. (2015). Arctic marine mammal population status, sea ice habitat loss, and conservation recommendations for the 21st century. Conservation Biology, 29(3), 724–737. 10.1111/cobi.12474

Laínez del Pozo, D., & Jones, P. J. (2021). Governance analysis of two historical MPAs in northern Peru: Isla Lobos de Tierra and Isla Lobos de Afuera. Marine Policy, 127, 104096. 10.1016/j.marpol.2020.104096

Lee, T., & McPhaden, M. J. (2010). Increasing intensity of El Niño in the central-equatorial Pacific. Geophysical Research Letters, 37(14). 10.1029/2010GL044007

Lehodey, P., Bertrand, A., Hobday, A. J., Kiyofuji, H., McClatchie, S., Menkès, C. E., … & Tommasi, D. (2020). ENSO impact on marine fisheries and ecosystems. El Niño Southern Oscillation in a changing climate, 429–451.

Lotze, H. K., & Worm, B. (2009). Historical baselines for large marine animals. Trends in ecology & evolution, 24(5), 254–262. 10.1016/j.tree.2008.12.004

Magera, A. M., Flemming, J. E. M., Kaschner, K., Christensen, L. B., & Lotze, H. K. (2013). Recovery Trends in Marine Mammal Populations. PLOS ONE, 8(10), e77908. 10.1371/journal.pone.0077908

Majluf, P., & Reyes, J. C. (1989). The marine mammals of Peru: a review. In The Peruvian upwelling ecosystem: dynamics and interactions. ICLARM Conference Proceedings (Vol. 18, No. 438, pp. 344–363).

McMahon, C. R., & Burton, H. R. (2005). Climate change and seal survival: Evidence for environmentally mediated changes in elephant seal, Mirounga leonina, pup survival. Proceedings of the Royal Society B: Biological Sciences, 272(1566), 923–928. 10.1098/rspb.2004.3038

Melin, S. R., Laake, J. L., DeLong, R. L., & Siniff, D. B. (2012). Age-specific recruitment and natality of California sea lions at San Miguel Island, California. Marine Mammal Science, 28(4), 751–776. 10.1111/j.1748-7692.2011.00538.x

Nicholls, N. (2008). Recent trends in the seasonal and temporal behaviour of the El Niño–Southern Oscillation. Geophysical Research Letters, 35(19). 10.1029/2008GL034499

Ouled-Cheikh, J., March, D., Borras-Chavez, R., Drago, M., Goebel, M. E., Fariña, J. M., Gazo, M., Coll, M., & Cardona, L. (2024). Future climate-induced distribution shifts in a sexually dimorphic key predator of the Southern Ocean. Global Change Biology, 30(3), e17191. 10.1111/gcb.17191

Peraltilla, S., Meneses, G., Gutiérrez, M., & Alfaro-Shigueto, J. (2024). The Salvamares Program of the National Fisheries Society: contributing to the sustainability of the Peruvian marine ecosystem. South Sustainability, 5(2), e102. 10.21142/SS-0502-2024-e102

Prentice, R. L., & Gloeckler, L. A. (1978). Regression analysis of grouped survival data with application to breast cancer data. Biometrics, 57–67. 10.2307/2529588

R Development Core Team. (2023). R: A language and environment for statistical computing (Version 4.4.1) [Computer software]. R Foundation for Statistical Computing. https://www.R-project.org/

Racault, M.-F., Sathyendranath, S., Brewin, R. J. W., Raitsos, D. E., Jackson, T., & Platt, T. (2017). Impact of El Niño Variability on Oceanic Phytoplankton. Frontiers in Marine Science, 4. 10.3389/fmars.2017.00133

Ramos, J. E., Tam, J., Aramayo, V., Briceño, F. A., Bandin, R., Buitron, B., Cuba, A., Fernandez, E., Flores-Valiente, J., Gomez, E., Jara, H. J., Ñiquen, M., Rujel, J., Salazar, C. M., Sanjinez, M., León, R. I., Nelson, M., Gutiérrez, D., & Pecl, G. T. (2022). Climate vulnerability assessment of key fishery resources in the Northern Humboldt Current System. Scientific Reports, 12(1), Article 1. 10.1038/s41598-022-08818-5

Rayner, N. A., Parker, D. E., Horton, E. B., Folland, C. K., Alexander, L. V., Rowell, D. P., Kent, E. C., & Kaplan, A. (2003). Global analyses of sea surface temperature, sea ice, and night marine air temperature since the late nineteenth century. Journal of Geophysical Research: Atmospheres, 108(D14), 4407. 10.1029/2002JD002670

Robertson, B. C., & Chilvers, B. L. (2011). The population decline of the New Zealand sea lion Phocarctos hookeri: A review of possible causes. Mammal Review, 41(4), 253–275. 10.1111/j.1365-2907.2011.00186.x

Sanderson, C. E., & Alexander, K. A. (2020). Unchartered waters: Climate change likely to intensify infectious disease outbreaks causing mass mortality events in marine mammals. Global Change Biology, 26(8), 4284–4301. 10.1111/gcb.15163

Schaub, M., & Abadi, F. (2011). Integrated population models: a novel analysis framework for deeper insights into population dynamics. Journal of Ornithology, 152(Suppl 1), 227–237.

Schaub, M., & Kéry, M. (2021). Integrated population models: Theory and ecological applications with R and JAGS. Academic Press. 10.1016/C2019-0-02015-8

Sepúlveda, M., Quiñones, R. A., Carrasco, P., & José Pérez-Álvarez, M. (2012). Daily and seasonal variation in the haul-out behavior of the South American sea lion. Mammalian Biology, 77(4), 288–292. 10.1016/j.mambio.2012.03.008

Sepúlveda, M., & Harcourt, R. G. (2021). Maternal behavior in otariids and the walrus. Ethology and behavioral ecology of otariids and the odobenid, 51–61. 10.1007/978-3-030-59184-7_3

Sepúlveda, M., Szteren, D., Alfaro-Shigueto, J., Crespo, E. A., Durán, L. R., Guerrero, A. I., … & Oliveira, L. R. (2023). Sea lion and fur seal interactions with fisheries and aquaculture in South American waters: threats and management perspectives. Mammal Review, 53(2), 116–131. 10.1111/mam.12311

Sepúlveda, M., Barilari, M. F., Deforest, K., Frantz, B., San Martin, M., Guerra, J., … & Hines, E. (2026). Bycatch risk assessment for South American sea lions using a GIS-based toolbox. Biological Conservation, 317, 111810. 10.1016/j.biocon.2026.111810

Sharples, R., Mackenzie, M., & Hammond, P. (2009). Estimating seasonal abundance of a central place forager using counts and telemetry data. Marine Ecology Progress Series, 378, 289–298. 10.3354/meps07827

Simmonds, M. P., & Isaac, S. J. (2007). The impacts of climate change on marine mammals: early signs of significant problems. Oryx, 41(1), 19–26. 10.1017/S0030605307001524

Soto, K. H. (2004). The effects of prey abundance on the diet, maternal attendance and pup mortality of the South American sea lion (Otaria flavescens) in Peru (Doctoral dissertation, University of British Columbia).

Soto, K. H., Trites, A. W., & Arias-Schreiber, M. (2006). Changes in diet and maternal attendance of South American sea lions indicate changes in the marine environment and prey abundance. Marine Ecology Progress Series, 312, 277–290. 10.3354/meps312277

Speakman, C. N., Hoskins, A. J., Hindell, M. A., Costa, D. P., Hartog, J. R., Hobday, A. J., & Arnould, J. P. Y. (2020). Environmental influences on foraging effort, success and efficiency in female Australian fur seals. Scientific Reports, 10(1), 17710. 10.1038/s41598-020-73579-y

Stan Development Team. (2023). Stan modeling language users guide and reference manual (Version 2.32.2) [Computer software]. https://mc-stan.org

Stoklosa, J., Blakey, R. V., & Hui, F. K. (2022). An overview of modern applications of negative binomial modelling in ecology and biodiversity. Diversity, 14(5), 320. 10.3390/d14050320

Stenson, G. B., & Hammill, M. O. (2014). Can ice breeding seals adapt to habitat loss in a time of climate change? ICES Journal of Marine Science, 71(7), 1977–1986. 10.1093/icesjms/fsu074

Taylor, M. H., Tam, J., Blaskovic, V., Espinoza, P., Ballón, R. M., Wosnitza-Mendo, C., … & Wolff, M. (2008). Trophic modeling of the Northern Humboldt Current Ecosystem, Part II: Elucidating ecosystem dynamics from 1995 to 2004 with a focus on the impact of ENSO. Progress in Oceanography, 79(2-4), 366–378.

Tinker, M. T., Carswell, L. P., Tomoleoni, J. A., Hatfield, B. B., Harris, M. D., Miller, M. A., Moriarty, M. E., Johnson, C. K., Young, C., Henkel, L. A., Staedler, M. M., Miles, A. K., & Yee, J. L. (2021). An integrated population model for southern sea otters (U.S. Geological Survey Open-File Report 2021–1076). U.S. Geological Survey. 10.3133/ofr20211076

Trites, A. W., & Larkin, P. A. (1996). Changes in the abundance of Steller sea lions(Eumetopias jubatus) in Alaska from 1956 to 1992: How many were there?. Aquatic Mammals, 22(3), 153–166.

Trivers, R. L. (1974). Parent-offspring conflict. American zoologist, 14(1), 249–264.

Vehtari, A., Gelman, A., & Gabry, J. (2024). loo: Efficient leave-one-out cross-validation and WAIC for Bayesian models (Version 2.8.0) [R package]. https://mc-stan.org/loo/

Wille, M., & Barr, I. G. (2022). Resurgence of avian influenza virus. Science, 376(6592), 459–460. 10.1126/science.abo1232

Wood, S. A., Josephson, E., Precoda, K., & Murray, K. T. (2022). Gray seal (Halichoerus grypus) pupping trends and 2021 population estimates in U.S. waters (NOAA Technical Memorandum NMFS-NE-314). U.S. Department of Commerce, Northeast Fisheries Science Center.

