## Supplemental Data 1 for "From Recovery to Risk: Intensifying ENSO Events Drive Extirpation Risk in a Marine Predator"

This word document containing detailed descriptions of:

SM1. Model structure and assumptions

SM2. Covariate sources and processing

SM3. Bayesian model implementation and evaluation

SM4. Projection framework

SD1. Compilation and harmonisation of count data from multiple sources

SR1. Model selection

SR2. Prior sensitivity analysis

Supplementary Methods

#### SM1. Model structure and assumptions

The population is assumed to be closed within Peruvian territory; genetic analyses indicate a close relationship between the Peruvian and northern Chilean populations (Oliveira et al., 2017), but long-term census data for northern Chile are limited, and telemetry studies indicate site fidelity across life stages (Baylis et al., 2017; Cárdenas-Alayza et al., 2022).

The model was structured as female-only, as incorporating males could increase the risk of overfitting given the complex model structure. Female-only models have commonly been employed in population models of pinnipeds and other taxa (Caswell, 2001; Schaub & Abadi, 2011), although we acknowledge that male dynamics can also influence overall population growth (Gerber & White, 2014).

To model the effect of SST, zooplankton concentrations, and anchovy biomass on the survival and birth probabilities, we used a complementary log-log (cloglog) regression with a quadratic term. We expect the covariate's influence on vital rates to intensify at extreme values, as *O. byronia* populations exhibit sharp declines during strong El Niño years (Arias-Schreiber & Rivas, 1998). The cloglog link is well-suited for capturing this pattern, as it models the log of the cumulative hazard and allows for an asymmetric response: probabilities can approach 0 more quickly than they approach 1 (or vice versa), depending on the direction of the effect. For example, survival may decline abruptly when prey availability drops below a critical threshold. Similarly, survival gains at higher prey levels may be minimal and plateau, resulting in a curved relationship better described by a quadratic form.

#### SM2. Covariate sources and processing

##### 2.1. Sea surface temperature

We used a simple spatial sampling framework to extract the dominant SST trend along the Peruvian coast. We selected three locations spanning northern, central, and southern latitudes to capture broad-scale spatial variation in SST. These regions do not represent strict ecological zones, but are positioned near the two ends of the population range and its midpoint. The SST dataset, developed by the Met Office Hadley Centre (https://hadobs.metoffice.com/), is available at a monthly resolution with a one-degree spatial grid and can be accessed via the Environmental Research Division's Data Access Program of the National Oceanic and Atmospheric Administration (NOAA) (<https://coastwatch.pfeg.noaa.gov/erddap/griddap/erdHadISST.html>). We used data spanning from April 1, 1995, to March 16, 2023, to perform a principal component analysis (PCA) and used the first principal component (PC1, which accounted for 95.72% of the total variance) as a proxy for the dominant SST trend along the coast. The sign of PC1 is arbitrary, and in this case, higher SST corresponds to lower PC1 values. We kept the original orientation, so that lower values consistently indicate higher environmental stress.

##### 2.2. Zooplankton concentrations

We used the same spatial sampling framework described above to extract the dominant zooplankton concentration trend along the Peruvian coast. We obtained monthly zooplankton carbon mass (zooc, g m⁻²) data from the Copernicus Marine Service dataset (cmems_mod_glo_bgc_my_0.083deg-lmtl_PT1D-i), which provides estimates from January 1, 1998, onwards. Since this dataset has finer spatial resolution than the SST data (0.083° grid), we initially selected three locations closer to major colonies that were positioned near the two ends of the population range and its midpoint. However, pairwise correlations among these three time series were low (Pearson’s r = 0.28–0.49, compared to 0.43–0.70 for SST), indicating high regional variability. To better capture this variation, we added four additional latitudinal points near major colonies distributed along the coast. We then performed PCA and used the PC1 (accounted for 46.62% of the total variance) as a proxy for the dominant zooplankton concentration trend along the coast. To extend the time series prior to 1998, PC1 values were hindcast using a monthly GAM with the SST proxy as the predictor. This approach resulted in a continuous monthly zooplankton anomaly time series covering the period from April 1, 1995, to March 16, 2023.

##### 2.3. Anchovy biomass

We extracted the anchovy biomass data from a stock assessment report (IMARPE, 2023). These data come from acoustic surveys that cover the 5°S to 15°S latitudinal range, which corresponds to the core distribution of the northern-central anchoveta stock. Data was digitized from the figure using WebPlotDigitizer (v3.4; Rohatgi, 2021), covering the years 1996 to 2023. Despite multiple requests, we were unable to obtain access to the full underlying dataset. As such, this digitized time series represents the best available approximation of stock biomass for this region. Two measurements are available each year, one in austral summer months (February to April) and one in the austral winter months (August to September).

For SST and zooplankton, the covariate time series are available at a monthly resolution. Therefore, we used the scaled mean of the monthly values within each temporal window (see Figure 2 in the main method) in the analysis. In contrast, anchovy biomass is reported seasonally, so we calculated the mean of seasonal values within each temporal window and scaled it accordingly. For example, to represent the 12 months before the breeding season in year $t$, the scaled mean of the austral summer in year $t$ and the austral winter in year $t-1$ was used.

#### SM3. Bayesian model implementation and evaluation

#### 3.1. Prior specification

##### 3.1.1. Vital rates

To complete the Bayesian model specification, we defined prior distributions for each model parameter based on available information. We defined priors for the stage-based survival rates and birth rate using life table data from an Atlantic population of *O. byronia* in Argentina (Grandi et al., 2016). We outlined the conversion process from age-specific to stage-specific survival rates in Chung et al. (2026). The life table data were derived from stranded animal records collected during a period of population recovery, similar to the situation for the Peruvian population during the study period. We defined scaled beta distributions centered on the literature-derived mean vital rates, with symmetric lower and upper bounds chosen to constrain parameters within biologically plausible limits, except for pup survival (Table SM1). Birth rate was parameterized as female pup production only, assuming an equal sex ratio at birth. For pup survival, we specified a wider upper range and an asymmetric beta distribution. This was intended to account for the expectation that environmental covariate effects would generally reduce realized survival values, thereby limiting the effective upper range available to the parameter with the lowest mean survival estimate among all stages (Table SM1). These bounds do not represent empirical confidence limits, but rather biologically reasonable constraints intended to stabilize estimation and avoid implausible parameter values (e.g., pup survival approaching 1). While demographic differences may exist between Atlantic and Pacific populations of the species, we assumed that the breadth of the prior distributions was sufficient to accommodate plausible variation in survival and reproductive rates across populations.

**Table SM1.** Prior distributions for all estimated parameters in the state-space model. Lower and upper bounds for vital rate baseline parameters ($\gamma_{r}$) define the biologically plausible range used to scale beta-distributed draws. Constraints on covariate ($\beta_{1,r}>0$, $\beta_{2,r}<0$) and process parameters ($\alpha_{r}> 0$,$\theta_{r} < 0$) reflect biological expectations of positive linear and negative quadratic environmental effects, and negative density dependence, respectively. Parameters marked $\dagger$ share a single prior draw across all vital rates $r$. Parameters marked $\ddagger$ share a single prior draw across juvenile age 1-3.

| Parameter | Description | Prior | Mean | Lower | Upper |
| --- | --- | --- | --- | --- | --- |
| $\gamma_{p}$ | Pup survival baseline | $Beta(1.49, 3.03)$ | 0.71 | 0.61 | 0.91 |
| ${\gamma_{j1}}^{\ddagger}$ | Age 1 juvenile survival baseline | $Beta(2, 2)$ | 0.88 | 0.78 | 0.98 |
| ${\gamma_{j2}}^{\ddagger}$ | Age 2 juvenile survival baseline | $Beta(2, 2)$ | 0.91 | 0.83 | 1.00 |
| ${\gamma_{j3}}^{\ddagger}$ | Age 3 juvenile survival baseline | $Beta(2, 2)$ | 0.94 | 0.88 | 1.00 |
| $\gamma_{f}$ | Adult survival baseline | $Beta(2, 2)$ | 0.93 | 0.86 | 1.00 |
| $\gamma_{br}$ | Birth rate baseline | $Beta(2, 2)$ | 0.30 | 0.10 | 0.49 |
| $h_{p}$ | Pup haul-out proportion | $Beta(5.39, 1.36)$ | 0.79 | 0 | 1 |
| $h_{f}$ | Female adult haul-out proportion | $Beta(10.87, 22.65)$ | 0.32 | 0 | 1 |
| $\beta_{1,r}$ | Linear covariate effect (all $r$) | $Cauchy(0, 0.1),$ $\beta_{1,r}>0$ | — | — | — |
| $\beta_{2,r}$ | Quadratic covariate effect (all  $r$) | $Cauchy(0, 0.1),$ $\beta_{2,r}<0$ | — | — | — |
| $\alpha_{r}$ | Environmental stochasticity SD (all  $r$) | $Cauchy(0, 0.1),$  $\alpha_{r}>0$ | — | — | — |
| $\theta_{r}$ | Density dependence coefficient (all  $r$) | $Cauchy(0, 0.1),$  $\theta_{r}<0$ | — | — | — |
| ${\varepsilon_{r,t}}^{\dagger}$ | Shared annual perturbation | $Normal(0, 1)$ | — | — | — |
| $\varphi_{1-3}$ | Negative binomial overdispersion (pup, female, total counts) | $Cauchy(0, 1),$  $\varphi>0$ | — | — | — |
| $\lambda₀$ | Initial population size (thousands) | $Cauchy(0, 1),$  $\lambda₀\geq150$ | — | — | — |
| $\delta$ | Sex ratio scaling factor (female-to-total) | $Normal(1, 0.5)$,  $\delta>0$ | — | — | — |
| $\varepsilon_{u,t}$ | Annual perturbation for scaling correction | $Normal(0, 1)$ | — | — | — |
| $\alpha_{u}$ | SD of scaling deviation | $Cauchy(0, 0.1),$  $\sigma_{u}>0$ | — | — | — |

##### 3.1.2 Haul-out ratios and correction factor

We estimated stage-specific haul-out ratios for adult females and pups to account for the fact that not all *O. byronia* are visible during surveys. Prior distributions for these ratios were informed by available data and their associated uncertainty. We used Beta distributions for both parameters to reflect haul-out probabilities bounded between 0 and 1. In addition, we included a multiplicative correction factor to account for years in which only total population counts (across all stages and sexes) were available. This factor was retained in all survey years to ensure a consistent model structure and to allow information about overall detectability and population composition to be shared across years. The prior specifications for the adult female haul-out ratio, pup haul-out ratio, and correction factor are described below.

The haul-out ratio of pups is defined as the proportion of pups present on a randomly selected sampling date relative to the highest number observed on a day across the survey period. This ratio was estimated using 12 years of weekly pup count data from Punta San Juan, a major breeding site (Data collected under: specific interinstitutional collaboration agreements between UPCH and SERNANP 2011-2016, 2017-2022 and RJ No. 013-2022-SERNANP-RNSIIPG). We assume that all pups born during the survey period are present onshore at the peak pupping date. A GAM was fitted to examine the relationship between this proportion and day of year within the survey period using a cubic spline basis (See Chung et al., 2026). The model explained 52.9% of the deviance, with smoothing parameter estimation performed using Generalized Cross-Validation (GCV). The smooth term for day of the year was significant (F = 12.78, p < 2 × 10⁻¹⁶). We use the mean and standard deviation (SD) of the predicted proportion to inform the shape parameter of the beta distribution for the prior of the pup haul-out ratio (Table SM1).

The haul-out ratio of adults is defined as the average proportion of adults present on a randomly selected sampling date. Unlike pups, adult females undertake long foraging trips during the breeding season (Soto et al., 2004), making it unlikely that any single count fully captures the total number present, regardless of the timing. Therefore, we used a separate Bayesian model to determine the mean and SD of the prior for the haul-out ratio of adults. We first specified a literature-informed prior (Romero et al., 2017), then updated it using telemetry data from Punta San Juan (RJ Nº 019-2016-SERNANP-RNSIIPG) in this separate Bayesian model. The only haul-out estimates available for this species in Romero et al. (2017) are not specified for adults and are likely to be too high for adults as pups are included, which have a high haul-out ratio. Therefore, we arbitrarily lowered the mean and chose a wide sd (beta ~ (2,2)) to make the prior less informative. Then, we used the proportion of time *O. byronia* were present in the rookery during daytime as the input data. We fitted this separate model using four Markov chains with 1,000 warm-up and 1,000 sampling iterations each, yielding 4,000 posterior samples (see Chung et al., 2026 for detailed specifications). We then used the resulting posterior distribution as the prior for the corresponding parameter in the population model (Table SM1).

For the correction factor, we assigned a weakly informative prior truncated to positive values, reflecting broad uncertainty while remaining consistent with biological expectations. The total population (both sexes) should exceed the female-only population but remain below twice that number, given sex-specific survival differences, e.g., higher male mortality through infanticide (Campagna et al., 1988). Detectability for hauled-out individuals, particularly males, is uncertain but highly unlikely to exceed that of pups (mean prior haul-out = 0.79). Centering the prior on 1 with a wide standard deviation of 0.5 therefore spans a plausible range of under- and over-correction (Table SM1). Because informative priors were already applied to the female and pup haul-out probabilities, this weak prior allows the correction factor to be informed by the data without unduly constraining the model.

##### 3.1.3. Model coefficients

To represent the impact of the covariate on vital rates in Equation 1, we included both linear and quadratic terms with corresponding coefficients, $\beta₁$ and $\beta₂$. We assigned half-Cauchy priors to these coefficients, following recommendations for regularisation of regression coefficients on bounded scales (Gelman et al., 2008; Table SM1). Exploring both positive and negative parameter spaces across all covariates and vital rates would be computationally inefficient and potentially increase the risk of overfitting. Therefore, we constrained the coefficients to reflect biologically plausible relationships. We retained the naturally inverted PC1 of SST that lower values correspond to higher temperatures during El Niño events. Thus, for all three covariates, lower values indicate increased hazard, reflecting the stronger population impact during El Niño when prey availability is reduced (Cárdenas-Alayza et al., 2021; de Oliveira et al., 2012). Using a complementary log-log link for survival modeling, we expected a steeper increase in hazard at the lower (negative) end of the covariate scale. This pattern was best captured by constraining the linear term ($\beta₁$) to negative values and the quadratic term ($\beta₂$) to positive values for all covariates, producing a concave downward response curve. This approach offers greater flexibility than a simple positive linear model while aligning with the biological expectation of stronger effects under extreme conditions. Similarly, $\theta$ parameters governing density dependence were restricted to negative values, aligning with expectations of negative density-dependent effects (Grandi et al., 2025; Romero et al., 2017).

#### 3.2. Model acceptance and assessment

##### 3.2.1. Acceptance criteria

To ensure efficient exploration of the posterior, we set the target acceptance rate to 0.99 and the maximum tree depth to 12. We considered a model to have converged if it met the following criteria: at least 99% of parameters had R-hat values below 1.01; bulk- and tail-effective sample sizes exceeded 100 per chain for 99% of parameters; more than 99% of post-warmup samples were not divergent transitions; no transitions reached the maximum tree depth; and all chains had energy-BFMI values greater than 0.3. We followed the convergence thresholds recommended by the Stan Development Team (2025), but manually adjusted the percentage thresholds (setting them at 99% rather than 100%) to avoid overly stringent requirements that may penalize otherwise well-behaved models.

##### 3.2.2. Model assessment

In the first stage, we calculated the total negative binomial log-likelihood for each posterior draw and applied LOO using the loo package (version 2.8.0). LOO provides an approximation of a model’s out-of-sample predictive accuracy, using Pareto-smoothed importance sampling to stabilize the estimates. The main diagnostic statistic is the expected log pointwise predictive density (elpd). We used the loo_compare() function to compute the difference in elpd between each model and the best-performing model (i.e., the one with the highest elpd), along with the standard error of that difference. If the 95% confidence interval of the difference (±1.96 SE) does not include zero, we interpret this as strong evidence that the best model has substantially better predictive performance than the model being compared. We also used Pareto k diagnostic values to quantify the reliability of this approximation. High Pareto k values (e.g., > 0.7) indicate influential observations that may cause instability or bias in the LOO estimates, suggesting that the LOO approximation might be less trustworthy for those points. Therefore, to complement the model ranking by LOO, we computed Fitted Bayesian R². Fitted Bayesian R² measures the proportion of variance in the observed data explained by the model’s fitted values. Following Gelman et al. (2019), it is calculated for each posterior draw as equation S1, where $\hat{y}$are the fitted values and $y-\hat{y}$ are the residuals. The result is a posterior distribution of R², summarised by its mean and credible intervals.

$R^{2}=Var(\hat{y})+Var(y-\hat{y})$	Equation [S1]

We applied LFO in the second stage, as LFO provides a more realistic measure of predictive performance than LOO, particularly for time-series data where observations are temporally autocorrelated. Unlike LOO, which removes a single observation at a time and allows information from the future to inform predictions of the past, LFO respects the temporal ordering of the data and avoids this type of information leakage (Bürkner et al., 2020). However, because LFO is computationally intensive and can be more difficult to achieve convergence, we applied it only to the ten best-performing models from stage one. In LFO, the data are split into a training set and a test set in a rolling-origin manner. We began with the first 10 years as the training set, as the population appeared to reach a stable recovery trend by year 10. We then fit the model to years 1–10 and predicted year 11, fit to years 1–11 and predicted year 12, and so on, skipping any blocks where the test year had no available data. To ensure that all blocks could be refitted for all models, we changed the random seed of some blocks and relaxed the percentage threshold of convergence to 2%. To aid comparison with the base model, for which LFO struggled to converge, we conducted posterior predictive checks for the top-scoring models and the base model. For each model, we extracted all posterior samples of the initial population size and parameters, excluding yearly estimates of population size and vital rates. We then simulated population trajectories by running the model forward using the corresponding covariate time series, while incorporating randomly generated environmental noise, correction factor noise, and observation error.

#### SM4. Projection

#### 4.1. Generation of covariate time series for projection

We used a three-step approach to generate covariate time series under the defined ENSO scenarios. First, we simulated ENSO state sequences using a Markov process based on historical transitions among three ENSO states, classified using the 12-month rolling average of the Multivariate ENSO Index (MEI) from April to March (1997–2022), spanning between the species’ breeding season. To simulate increased frequency, we adjusted transition probabilities so that El Niño and La Niña years increased by 30% (medium) and 59% (high), respectively. Second, we modelled ENSO-driven changes in covariate anomalies by estimating the historical MEI distribution using kernel density estimation (KDE). We generated medium and high intensity scenarios by scaling the spread of MEI values by ±10% and ±20% around the mean. Third, we translated MEI values into covariate anomalies using a two-step GAM: one for the MEI-dependent mean and another for the MEI-dependent standard deviation, based on absolute residuals. This allowed us to sample realistic future covariate time series from a normal distribution that reflects both observed mean and variability.

#### 4.2. Extirpation risk metrics

We applied the IUCN Red List A2 criterion (IUCN, 2012) to estimate the probability of the population crossing the critically endangered threshold, defined as a ≥80% decline over three generations. Based on previous studies on otariids (Weinberger et al., 2021; Beauplet et al., 2006; Boyd et al., 1999; Grau & Acuña, 1998), we assumed a generation time of 10 years. Therefore, any simulation in which the population declined by ≥80% within any rolling 30-year window over the 50-year projection period was recorded as meeting the IUCN Criterion A2 Critically Endangered threshold. The proportion of such simulations was calculated for each scenario, representing the likelihood of reaching this conservation threshold.

The extirpation outcome was binary (crossing the critically endangered threshold or not), thus we calculated 95% confidence intervals (CI) for each estimated probability using Wilson score intervals. The Wilson Score confidence intervals reflect uncertainty in the estimated extirpation proportion conditional on the realized simulation ensemble, but do not capture additional variability associated with rerunning the stochastic projection process. Each scenario included 11,998 simulation replicates, which encompassed both parameter uncertainty (from the full posterior distribution of the fitted population model) and stochastic variability in future ENSO conditions. As such, the resulting extirpation risk estimates and CIs incorporate both model and environmental uncertainty.

**Table SM2.** Prior sensitivity analyses applied to the highest-scoring model. Eleven alternative prior specifications were evaluated independently to assess sensitivity to prior assumptions. Variations were grouped as global (affecting all vital rates or haul-out parameters simultaneously) or local (targeting specific demographic processes).

| Category | Sensitivity variation | Variables affected | Extent |
| --- | --- | --- | --- |
| Global | Flatten the prior distribution | All vital rates | 0.5 |
| Global | Left shift of the prior distribution | All vital rates | 0.05 |
| Global | Narrow the prior distributions | Pup and adult haul-out ratios | 0.5 |
| Local | Left shift of the prior distribution | Pup survival rate | 0.05 |
| Local | Left shift of the prior distribution | All juvenile survival rates | 0.05 |
| Local | Left shift of the prior distribution | Adult survival rate | 0.05 |
| Local | Left shift of the prior distribution | Birth rate | 0.05 |
| Local | Flatten the prior distribution | Pup haul-out ratio | 0.5 |
| Local | Flatten the prior distribution | Adult haul-out ratio | 0.5 |
| Local | Narrow the prior distribution | Pup haul-out ratio | 0.5 |
| Local | Narrow the prior distribution | Adult haul-out ratio | 0.5 |

Note: Adjustments consisted of either a proportional decrease of 0.05 in the prior mean (left shift) or a proportional modification of 0.5 in prior dispersion, with narrowing representing increased prior certainty and flattening representing decreased prior certainty.

References

Arias-Schreiber, M., & Rivas, C. (1998). Distribución, tamaño y estructura de las poblaciones de lobos marinos arctocephalus australis y otaria byronia en el litoral peruano, en noviembre 1996 y marzo 1997 (No. 73). Instituto del Mar del Perú.

Baylis, A. M., Orben, R. A., Costa, D. P., Tierney, M., Brickle, P., & Staniland, I. J. (2017). Habitat use and spatial fidelity of male South American sea lions during the nonbreeding period. Ecology and evolution, 7(11), 3992-4002. <https://doi.org/10.1002/ece3.2972>

Beauplet, G., Barbraud, C., Dabin, W., Kussener, C., & Guinet, C. (2006). Age-specific survival and reproductive performances in fur seals: Evidence of senescence and individual quality. Oikos, 112, 430–441. <https://doi.org/10.1111/j.0030-1299.2006.14412.x>

Boyd, I. L., Lockyer, C., & Marsh, H. D. (1999). Reproduction in marine mammals. In J. E. Reynolds, & S. A. Rommel (Eds.), Biology of marine mammals (pp. 218–286). Smithsonian Institution.

Bürkner, P. C., Gabry, J., & Vehtari, A. (2020). Approximate leave-future-out cross-validation for Bayesian time series models. Journal of Statistical Computation and Simulation, 90(14), 2499-2523.

Campagna C., & Boeuf B. J. L. (1988). *Reproductive Behaviour of Southern Sea Lions*. <https://doi.org/10.1163/156853988X00539>

Cárdenas-Alayza, S., Gutiérrez, D., & Tremblay, Y. (2021). Trends in sympatric otariid populations suggest resource limitations in the Peruvian Humboldt Current System. *Marine Environmental Research*, *169*, 105349. <https://doi.org/10.1016/j.marenvres.2021.105349>

Cárdenas-Alayza, S., Adkesson, M. J., Gutiérrez, D., Demarcq, H., & Tremblay, Y. (2022). Strategies for segregation during foraging in sympatric otariids of the Peruvian upwelling Humboldt Current System. Marine Ecology Progress Series, 702, 153-170. <https://doi.org/10.3354/meps14203>

Caswell, H. (2001). Matrix population models: construction, analysis, and interpretation. In Matrix population models: construction, analysis, and interpretation (pp. 722-722).

Chung, H. W. Y., Cárdenas-Alayza, S., Alfaro Shigueto, J., Farfán-Hernandez, K., Ganoza-Gallardo, M. J., Mengel, J. C., Davis, K., & Bonsall, M. (2026). Supplementary Code and Workflows for "From Recovery to Risk: Intensifying ENSO Events Drive Extirpation Risk in a Marine Predator". Zenodo. https://doi.org/10.5281/zenodo.20394833

De Oliveira, L. R., Fraga, L. D., & Majluf, P. (2012). Effective population size for South American sea lions along the Peruvian coast: The survivors of the strongest El Niño event in history. *Journal of the Marine Biological Association of the United Kingdom*, *92*(8), 1835–1841. https://doi.org/10.1017/S0025315411001871

Gelman, A., Jakulin, A., Pittau, M. G., & Su, Y.-S. (2008). A weakly informative default prior distribution for logistic and other regression models. *The Annals of Applied Statistics*, *2*(4), 1360–1383. <https://doi.org/10.1214/08-AOAS191>

Gelman, A., Goodrich, B., Gabry, J., & Vehtari, A. (2019). R-squared for Bayesian regression models. The American Statistician.

Gerber, L. R., & White, E. R. (2014). Two‐sex matrix models in assessing population viability: when do male dynamics matter?. Journal of Applied Ecology, 51(1), 270-278.

Grandi, M. F., Dans, S. L., & Crespo, E. A. (2016). Improvement in Survivorship: The Key for Population Recovery? *Zoological Studies*, *55*, e9. https://doi.org/10.6620/ZS.2016.55-9

Grandi, M. F., Ayelén, T., Reyes, L. M., García, N. A., & Crespo, E. A. (2025). Tracking South American Sea Lion Population Recovery in Patagonia Following Depletion From Hunting. Marine Mammal Science, 41(4), e70041. <https://doi.org/10.1111/mms.70041>

Grau, R., & Acuña, E. (1998). Determinación de la edad y crecimiento en el lobo marino común Otaria flavescens (Shaw 1800), a través del análisis de estructuras dentales por corte, en la zona norte de Chile. XVII Congreso de Ciencias del Mar, Resúmenes. (pp. 95–96).

IMARPE. (2023). Informe sobre la situación del stock norte-centro de la anchoveta peruana (Engraulis ringens) al 25 de mayo y perspectivas de explotación para la primera temporada de pesca de 2023. Instituto del Mar del Perú. Retrieved from [<https://www.gob.pe/institucion/imarpe/informes-publicaciones/4283395-informe-sobre-la-situacion-del-stock-norte-centro-de-anchoveta-peruana-engraulis-ringens>]

IUCN. (2012). IUCN Red List categories and criteria: Version 3.1 (2nd ed.). IUCN. <https://portals.iucn.org/library/node/10315>

Oliveira, L. R. D., Gehara, M. C., Fraga, L. D., Lopes, F., Túnez, J. I., Cassini, M. H., ... & Bonatto, S. L. (2017). Ancient female philopatry, asymmetric male gene flow, and synchronous population expansion support the influence of climatic oscillations on the evolution of South American sea lion (Otaria flavescens). PloS one, 12(6), e0179442. <https://doi.org/10.1371/journal.pone.0179442>

Rohatgi, A. (2021). WebPlotDigitizer (Version 3.4) [Computer software]. <https://automeris.io/WebPlotDigitizer>

Romero, M. A., Grandi, M. F., Koen-Alonso, M., Svendsen, G., Ocampo Reinaldo, M., Garcia, N. A., ... & Crespo, E. A. (2017). Analysing the natural population growth of a large marine mammal after a depletive harvest. Scientific Reports, 7(1), 5271.

Schaub, M., & Abadi, F. (2011). Integrated population models: a novel analysis framework for deeper insights into population dynamics. Journal of Ornithology, 152(Suppl 1), 227-237.

Soto, K. H. (2004). The effects of prey abundance on the diet, maternal attendance and pup mortality of the South American sea lion (Otaria flavescens) in Peru (Doctoral dissertation, University of British Columbia).

Weinberger, C. S., Vianna, J. A., Faugeron, S., & Marquet, P. A. (2021). Inferring the impact of past climate changes and hunting on the South American sea lion. *Diversity and Distributions*, *27*(12), 2479–2497. https://doi.org/10.1111/ddi.13421

Supplementary Data 1 - Combining Count Data from Multiple Sources

To compile a complete dataset of South American Sea Lion (SASL) population total counts and stage counts (only counts for pup and adult female are used in this model), we used three main sources:

1. Total Abundance Data (2000–2019): Obtained through a request for information through the Ley de Transparencia to IMARPE in 2020, referred to as "letter 1" (N° 149-2020).

2. Stage-Based Count Data (2000–2018): Also obtained from IMARPE in 2020. This dataset lacks pup counts and is referred to as "letter 2" (N° 070 -2020-IMARPE/OGA).

3. IMARPE Annual Reports (1997–2022): These reports include total abundance for most years and stage-based counts for selected years, referred to as the "reports".

Despite originating from the same institution and surveys, discrepancies were observed between the data sources, particularly between "letter 1" and the reports, and between "letter 2" and the reports. However, the data were generally consistent. Table 1 outlines the reasons behind our choice of data sources for each year, as well as the rationale for excluding certain years.

After creating a tidy dataset, we examined the correlation between pup and female count data with the total population count. Both the pup and female counts showed a significant correlation with the total count (P < 0.01). However, as indicated by the diagnostic plots in Figure SD1, one adult female count was identified as an outlier and subsequently excluded from further analysis.

**Table SD1.** Data sources of total count and stage-based count

| **Year** | **Total count** | **Stage-based count** | **Remarks** |
| --- | --- | --- | --- |
| 1997 | report | NA | Data is only available in the corresponding source |
| 1998 | report | NA | Data is only available in the corresponding source |
| 1999 | report | NA | Data is only available in the corresponding source |
| 2000 | report | report | Total count in the report is 14% higher, but it is preferred as the explanation in the paragraphs is detailed, and both pup and female adult counts are available |
| 2001 | NA | NA | No data |
| 2002 | report | letter 2 & report * | The percentage difference is less than 7% |
| 2003 | report | letter 2 & report * | The percentage difference is less than 7% |
| 2004 | report | letter 2 & report * | The percentage difference is less than 7% |
| 2005 | report | letter 2 & report * | The percentage difference is less than 7% |
| 2006 | report | letter 2 & report * | The percentage difference is less than 7% |
| 2007 | report | report | No discrepancy in total count, minimal discrepancy in female adult count |
| 2008 | report | report | No discrepancy in total count, minimal discrepancy in female adult count |
| 2009 | letter 1 | letter 2 | Total count in letter 1 is 11% higher, but it is preferred as the female adult count is only available in letter 2 |
| 2010 | discarded | discarded | The report indicates a three-week delay. Only the female adult count is available, so we discarded this year. |
| 2011 | discarded | report | The report indicates a three-week delay. Pup count is corrected with the same approach of determining the pup haul-out ratio^ |
| 2012 | discarded | discarded | Some important breeding sites were not visited (e.g. the Paracas Reserve) |
| 2013 | discarded | discarded | The sum of stage-based counts (excluding pup) is only 30% of the total count |
| 2014 | discarded | discarded | High discrepancy (>60%) |
| 2015 | report | report | Total count is 10% lower, and female count is 5% lower in the report, but it is preferred as pup count is only available in the report |
| 2016 | report | report | No discrepancy in total count, minimal discrepancy in female adult count |
| 2017 | report | letter 2 & report * | No discrepancy in total count |
| 2018 | report | letter 2 & report * | No discrepancy in total count |
| 2019 | report | report | No discrepancy in total count |
| 2020 | report | report | Data is only available in the corresponding source |
| 2021 | discarded | discarded | The survey was significantly delayed (2 months) |
| 2022 | report | report | Data is only available in the corresponding source. The report indicated that fewer colonies were surveyed due to a budget cut, but the most important breeding colonies where most pups were born were included. Therefore, the pup count was retained for this year. |

* Pup count from report, as letter 2 does not include pup count.

^ Code available at Chung et al. (2026).

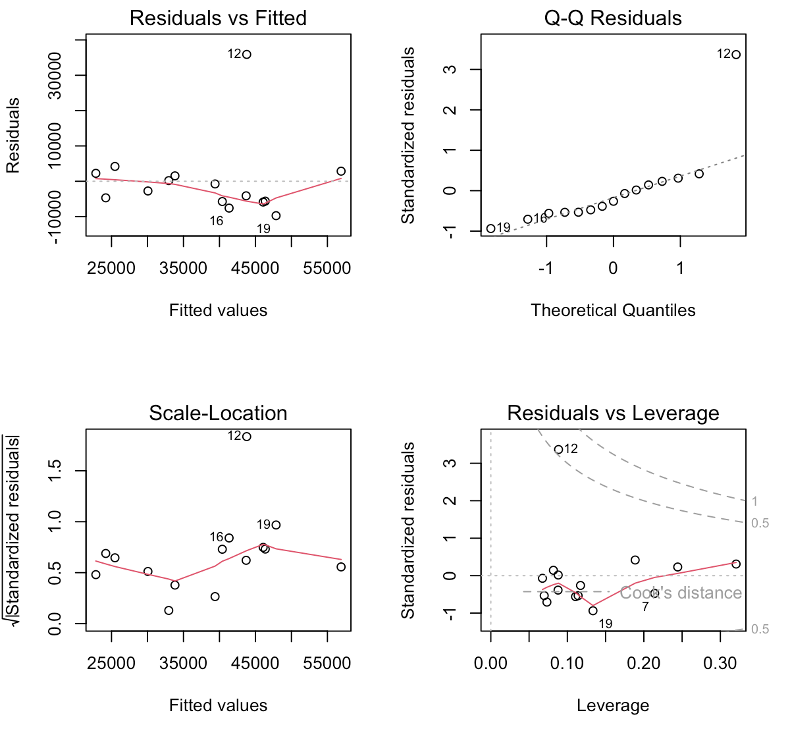

**Figure SD1.** Diagnostic plots showing the correlation between female count and total population count, datapoint 12 is identified as an outlier.

Reference

Chung, H. W. Y., Cárdenas-Alayza, S., Alfaro Shigueto, J., Farfán-Hernandez, K., Ganoza-Gallardo, M. J., Mengel, J. C., Davis, K., & Bonsall, M. (2026). Supplementary Code and Workflows for "From Recovery to Risk: Intensifying ENSO Events Drive Extirpation Risk in a Marine Predator". Zenodo. <https://doi.org/10.5281/zenodo.20394833>

### Supplementary Results 1&2

#### SR1. Model selection

Eleven of the 59 covariate models failed to meet the convergence thresholds, even after three attempts, and were excluded from further analysis. In the first stage of analysis, both LOO and fitted R² indicated that the ten highest-performing models all used survival covariates from the 12 months preceding the breeding season for five life stages. Within this top group, the best two models, one incorporating zooplankton and one incorporating SST anomalies, both apply the same one-year lag to survivals and birth rate. The top model was not significantly better than the other leading models or the base model, but it outperformed the remaining 37 models by a statistically significant margin in LOO, as indicated by the 95% CI of the elpd difference overlapping zero (Figure SR1). All LOO models, however, had between 4.2% and 37.5% of Pareto k values above 0.7, suggesting possible bias in the PSIS estimates (see Discussion). R² confidence intervals were wide, so we could not distinguish a significantly better model within the leading group. However, the top ten models according to R² were consistent with LOO in identifying models with biologically coherent covariate and lag structures. Based on this agreement, we proceeded to the second stage using these ten models plus the base model.

In the second stage, we evaluated this group of models using LFO cross-validation and posterior predictive checks. LFO is more appropriate for time-series evaluation because it tests the model’s ability to predict future observations, using fully independent refits for each training block to provide exact one-step-ahead predictions. LFO is, however, computationally intensive and prone to convergence issues; one covariate model and the base model failed to fully converge after three attempts. Among the models that did converge, the best SST-based model from stage 1 (model 19 in figure SR1) achieved the highest elpd, slightly outperforming the zooplankton group. Even the best-performing zooplankton model from stage 1 had a lower LFO elpd (−293.96 ± 4.76) compared with this SST model (−292.46 ± 4.50), corresponding to an elpd difference of −1.50 ± 1.20. Although standard errors of LFO were wide, the overall ranking supports the choice of SST for its better time-series predictive performance. This model was also in a close tie for the top two in the first stage with the zooplankton model. The PPC plots (Figure SR2) further highlight its ability to reproduce observed population fluctuations compared with the base model.

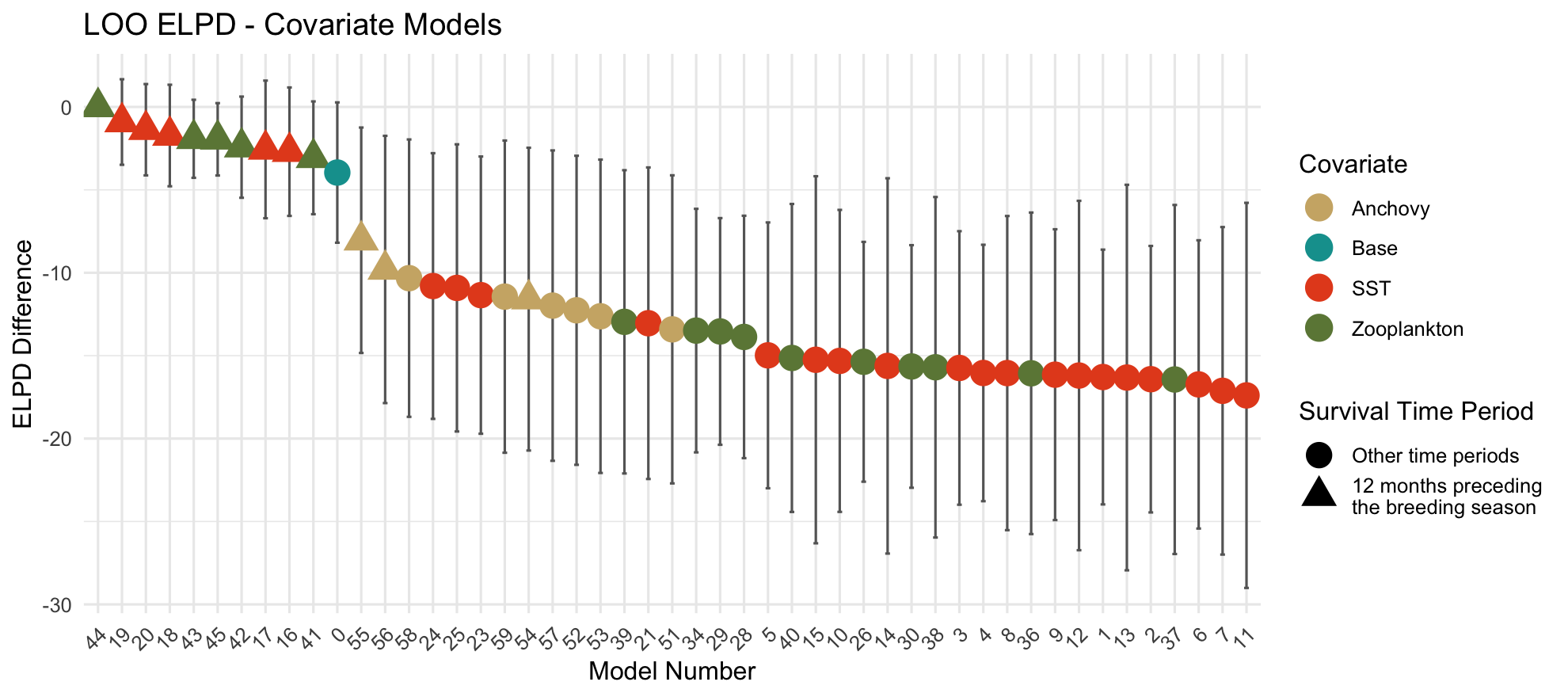

**Figure SR1.** Expected LOO ELPD differences between models relative to the best-performing model, ranked from highest to lowest. Error bars represent 95% CIs of the differences.

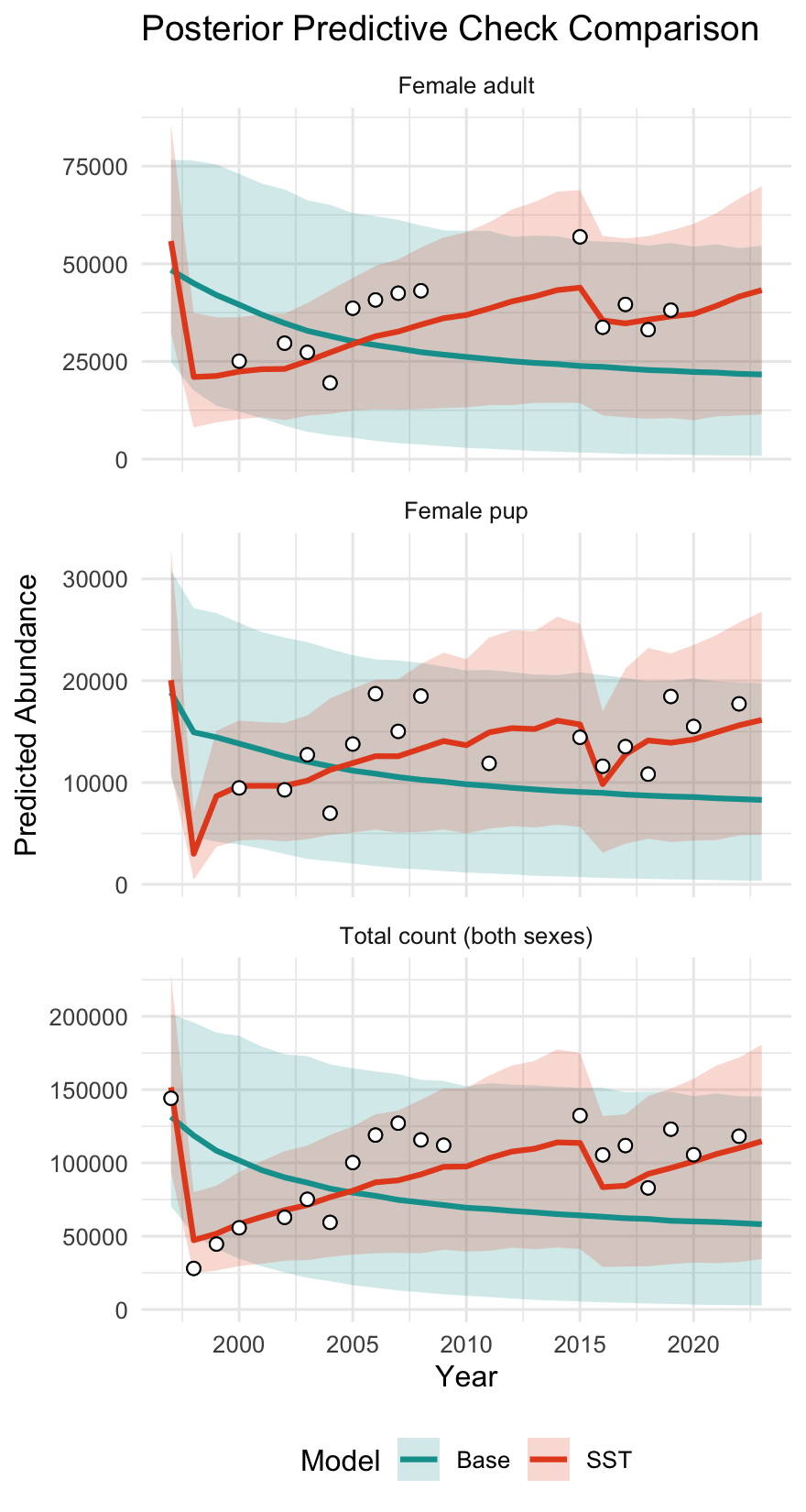

**Figure SR2.** Posterior predictive check comparing observed counts (black-outlined white points) to model-predicted abundance (lines and shaded ribbons) for the best-performing model in LFO analysis (SST model) and the base model, across female adults, female pups, and total abundance. Shaded areas represent 90% credible intervals. The monotonic decline of the base model reflects the deterministic effect of density dependence acting on a large initial population without environmental recovery signals.

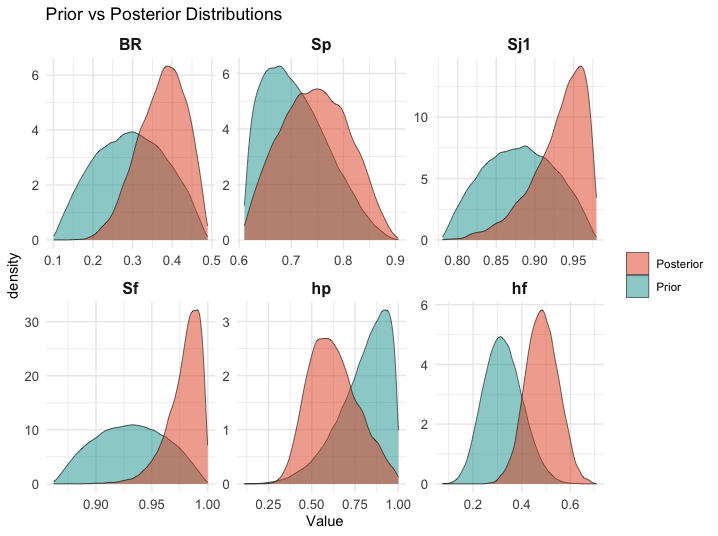
**Figure SR3.** Prior–posterior overlap for key demographic parameters: total birth rate of both sexes (BR), pup survival (Sp), juvenile survival in year 1 (Sj1), subadult female survival (Sf), and haul-out ratios of pups (hp) and adult females (hf).

#### SR2. Prior sensitivity analysis

Population fluctuations over the years were largely consistent across scenarios, with all trends capturing the drastic declines during the two major El Niño events. To compare each time series with the original, we calculated the absolute percent change in total female population estimates for each year and then averaged these values across years. The resulting means were ≤6.3% in all scenarios, except for the narrowing female haul-out scenario, which showed a substantially higher mean of 11.6% and a pronounced upward shift in the time series (Figure SR4). In contrast, flattening the adult haul-out and narrowing the pup haul-out led to the largest downward shifts.

LOO elpd values were not significantly different from the original model in most scenarios, except for flattening pup haul-out, flattening adult haul-out, and shifting adult survival left, where the confidence intervals showed slight non-overlap with the original (Figure SR5). These results should be interpreted with caution, however, as 25.0–37.5% of the LOO Pareto k values exceeded 0.7. The flexibility of the model, including the environmental stochasticity term, likely explains the highly similar trajectories and thus the similar LOO elpd values across scenarios.

Posterior estimates of baseline vital rates across the sensitivity scenarios were also similar to those of the original model, as the covariate and environmental effects largely determine the actual yearly vital rate estimates. Absolute percentage changes in baseline survivals were ≤4.91% across all stages and scenarios (mean = 1.02%). Birth rate showed greater changes, for example, decreasing by 8.45% when both haul-out ratios were narrowed. Haul-out ratios changed the most, as expected when their priors were modified. Pup haul-out increased by 10.66% and adult haul-out decreased by 12.68% when their prior distributions were narrowed by 50% respectively.

Compared to the above metrics, covariates, environmental stochasticity, and density-dependent effects were much more sensitive to prior changes. Density-dependent coefficients changed the most when the priors of all vital rates were decreased by 5%, with shifts ranging from −68.02% to −24.27%. The increases and decreases of the other two effects across stages were less consistent. We therefore ranked the sensitivity of each scenario using the sum of absolute changes. For the covariate effect, we focused on the first-order coefficient in the quadratic equation, as it indicates how strongly El Niño conditions reduced survival. For both the covariate and environmental stochasticity coefficients, decreasing the adult survival prior by 5% produced the most sensitive scenario. Changes across stages for the covariate and environmental stochasticity coefficients ranged from −41.65% to 6.14% and −24.14% to 29.03%, respectively.

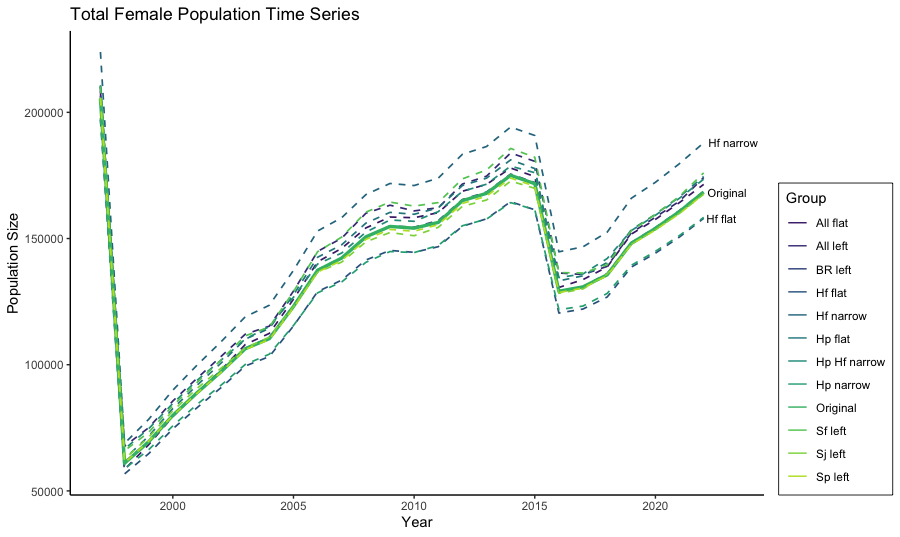

**Figure SR4.** Time series of population trajectories generated under different sensitivity scenarios. The solid line represents the original time series, while dashed lines indicate outcomes under alternative scenario settings. The lowest line corresponds to flattening adult haul-out, followed by narrowing pup haul-out. For scenario abbreviations used in this figure, please refer to **Table SM2**.

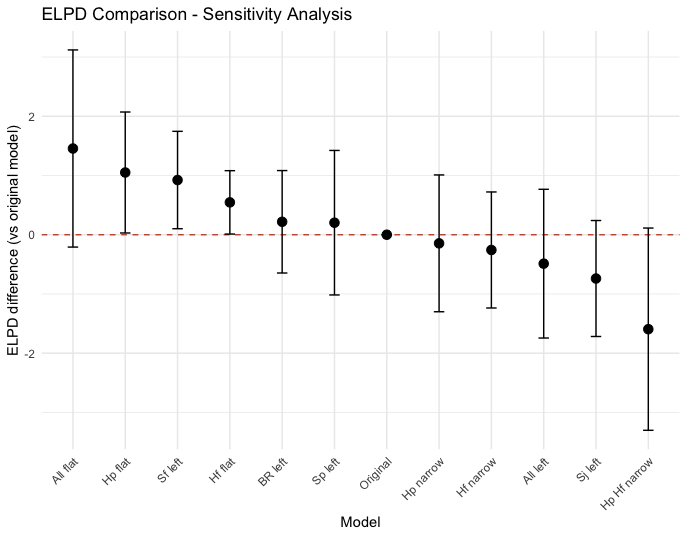

**Figure SR5.** ELPD difference of model variations evaluated in the sensitivity analysis. Error bars indicate 95% confidence intervals of the ELPD difference relative to the original model. See **Table SM2** for detailed scenario descriptions.
